# The Naïve Bayes Classifier++ for Metagenomic Taxonomic Classification – Query Evaluation

**DOI:** 10.1101/2024.06.25.600711

**Authors:** Haozhe (Neil) Duan, Gavin Hearne, Robi Polikar, Gail Rosen

## Abstract

This study examines the query performance of the NBC++ (Incremental Naive Bayes Classifier) program for variations in canonicality, kmer size, databases, and input sample data size. NBC++ can successfully assess a wide range of superkingdoms using a small training database. We demonstrate that NBC++ and Kraken2 are affected by database depth with macro measures increasing with depth but that the full diversity of life, especially viruses, is still a challenge for these classifiers. NBC++ spends less time training but at the cost of long querying time. The major enhancements are to accommodate canonical *k*mer storage (with major storage savings), adaptable and optimized memory allocation that quickens the query analysis and allows the classifier to be run on almost any system, and enables output of the log-likelihood values against each training genome which provides users with valualbe confidence information.

## 1 Introduction

The näive Bayes classifier (NBC) has proven itself to be a useful tool for classifying reads from amplicon and metagenomic samples [9, 6, 7, 2, 11]. For 16S rRNA taxonomic classification, it is the gold standard [13] due to its superior ability to memorize kmers unique to different taxa. For metagenomics, even with a limited database, it was very successful in determining the most relevant taxa and their abundances in most samples [4].

While NBC is computationally tractable for 16S rRNA due to its simplicity, NBC has not expanded for its use for metagenomics because of scalability issues, as well as not being able to easily provide a confidence for its prediction (without adding significant computational time). There are two specific scalability issues:1) One issues is that *k*mers must be counted during training, which is time consuming, and their frequencies must be calculated [11]. In [11], we addressed this issue, by adding a capability of adding genomes to the database without having to recompute all *k*mer frequencies of the entire database again. 2) Another issue is in the testing process of evaluating queries. In the previous implementation of NBC++ [11], *k*mer counting of the queries was conducted utilizing Jellyfish [3] and classess were added incrementally to the database. Due to Jellyfish requiring a file for each separate count, this approach often resulted in the creation of an extensive number of files, frequently reaching hundreds of thousands or even millions for a single metagenomic file. To mitigate this challenge, the current study introduces a novel runtime counting mechanism within the testing module to reduce disk I/O.

We examined a variety of parameters that offer speedups (sometimes at the cost of recall/precision), such as *k*mer size and canonical vs. non-canonical kmer counting. We also experiment with three different database sizes to show the speed vs. accuracy trade-off of using small but diverse datasets vs. large, rich databases when compared to Kraken2. Finally, we evaluate how long it takes for NBC++ to classify a large-scale real human metagenomic sample when trained on increasing database depths.

## 2 Improvements and Methodology since Zhao et al. [11]

The aim of our previous research was to augment the size of the training database and enable incremental database updates. In this paper, we depart from the training side and introduce a suite of enhancements aimed at refining the querying mechanism. These improvements are articulated as follows:

1. We optimized memory allocation within the confines of existing memory limitations, thus ensuring a more effective deployment of computational resources and adherence to memory constraints.
2. We introduce new features that facilitate the generation of full log-likelihood outputs, enabling a more thorough examination of the result distribution by users.
3. We implement canonical counting techniques which has led to a notable reduction in both the volume of training data and the disk space required for storing kmer counts, bolstering the system’s overall efficiency.
4. We restructure the querying operational protocol to decrease frequent loading of training data from disk (when possible - the user must specify a relatively large memory limit), which in turn mitigates computational overhead and enhances the efficiency of the querying process. A severe limit of the previous implementation required that the reads be pre- *k*mer counted before the query NBC++ code and that each read’s *k*mer count be stored in a separate file creating a large disk I/O load. This has now been restructured to count reads on the fly and to use the user-specified memory to determine how many to compute in a batch.

### 2.0.1 Allocation of Memory

The querying part of NBC++ has been improved to be able to fit into memory specified by the user, and can adapt to almost any memory size (a minimum of 2GB is suggested). This is in contrast to Kraken2, which takes up to 9GB for processing the Basic database, 107GB processing for the Standard database, and 208GB for the Extended database. Most of our experiments were performed with 192GB of memory; however, we do a 64GB memory comparison (as a desktop example).

### 2.0.2 Setting Capacities

In the NBC++ system, each processing thread is allocated:

- a workload capacity of 1000 query reads.
- an output buffer capacity of 1000 entries.

These capacities are the result of optimization on Illumina simulated data for minimizing read and write delays.

### 2.0.3 Estimating Memory Costs

Memory costs within the NBC system are estimated based on:

- the average memory size of a trained class. NBC++ querying memory consumption depends heavily on the *k*mer size used and the length of the genome trained on. For example Phytophthora infestans’s 9mer file is 1.8MB and its 15mer file is 818 MB, so 15mers requires approximately 400 × more memory at runtime.
- The average memory size per query read. Usually query reads are small (∼ 100-200bp for Illumina), but occasionally, researchers may want to use contigs or even full genomes.

See the Supplementary Material Appendix 1 for equations on how the input and output buffers of the querying process are reserved and memory is utilized.

### 2.0.4 Log-likelihood score outputs

There are now two modes of log-likelihood outputs. The default output produces the **max** log-likelihood score and the taxa corresponding to the **max** log-likelihood score, followed by the lineage of that taxa (upper taxonomic levels). We also implemented a feature so that users can explore the log-likelihood values of each query against each genome in the database, with the `-f` flag. This verbose output will increase the output tabular file greatly, as you will have each query’s log-likelihood score against each genome in the database and not just the max. You will likely need to adjust the `-r` and `-c` options to output reasonably sized tabular outputs. (The defaults are approximately the limits of what Excel can ingest).

#### 2.1 Data Collection and Database Construction

This study uses the Refseq retrieval pipeline developed as a part of the Woltka software [12] to sample the NCBI RefSeq database at high microbiome diversity at varying degrees of depth, and then test the performance of NBC++ on the resulting databases.

### 2.1.1 Database Construction

We used Woltka’s `refseq_build.py` script to extract genomic data from the RefSeq database. This script was employed to create databases with distinct profiles:

- “Basic” database: Comprising one genome per genus (with exceptions of Baxterfoxvirus and Betanucleorhabdovirus) resulting in a compilation of 4,634 genomes as of July 24, 2023.
- “Standard” database: Encompassing all NCBI-defined reference and representative genomes, totaling 18,237 genomes collected on July 26, 2023.
- “Extended” database: Featuring one genome per species with a Latinate name and higher ranks, accumulating 319,554 genomes by July 26, 2023. The extended database also includes reference, representative, and type material genomes. The genomes were then grouped by species resulting in: **4**,**634 classes** for “Basic,” **18**,**219 classes** for “Standard,” and **58**,**978 classes** for “Extended.” (see supplementary assembly files*_assemblysummary.txt, where * is basic/standard/extended)

### 2.1.2 Data Sources and Experimental design

#### Simulated Data

For all simulated experiments, we used a 5-fold cross-validation (CV) framework. First, we shuffled the dataset. Subsequently, we set the first one-fifth segment of this shuffled array as the testing set for the first fold, with the remaining data serving as the corresponding training set. We repeated this procedure five times to generate the 5 folds. For the generation of testing reads, we used InSilicoSeq [1] to synthesize 100 reads per class, utilizing the metagenomic file that was sampled from the RefSeq database.

#### Real Human Gut Metagenomic Sample

To assess a real sample, we again use the data from human gut sample assessed in [5, 11]. With such a sample, we can assess the composition of the sample with relative abundances (RAs) and compare against various parameters with the Bray-Curtis distance [11].

## 3 Results

We benchmark *k*mer size, canonicality, and database depth on the querying results and time. For training, the number of *k*mers is counted and either stored canonically or non-canonically. Since add-1 (Laplacian) smoothing is used for estimating the probability of the *i*_*th*_ *k*mer in a genome as 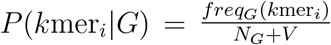 where freq_*G*_(*k*mer_*i*_) is the *i*th *k*mer in genome *G, N*_*G*_ is the number of *k*mers in genome *G*, and *V* is the vocabulary size of all *k*mers in all genomes. *V* can be approximated at *V* = 4^*k*^ whenever a *k*mer occurs at least once in one of the genomes in the entire database. For canonical *k*mers, for *k*mers of even length, the formula becomes 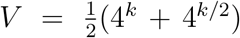, while for those of odd length, it simplifies to 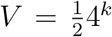 [10]. These estimates are then used in the maximum likelihood formulation of NBC [6].

### 3.1 Parameter Sweeps

#### 3.1.1 Canonical vs. Non-canonical

In the present update, we switched to employing canonical representations for the enumeration and categorization of sequences, thereby achieving a reduction in computational time and memory allocation for resultant savefiles. Canonical counting chooses between the *k*-mer in question and its reverse complement, with the lexicographically smaller sequence being selected for utilization. Conversely, non-canonical counting involves the direct application of the *k*-mer as it stands. As illustrated in Supplementary Fig. 1, canonical and non-canonical counting exhibit comparable efficacy in the context of Basic 9-mer and Standard 9-mer. While there is little querying time speedup with the canonical representation, the amount of disk space saved can vary. For example, at 9mers, where almost every 9mer exists in long genomes, the canonical count representation uses 54% of the disk space (compared to the noncanonical representation). For 15mers, the disk savings diminish – with long microbes still taking up 90% and the longest fungi taking 83% of the noncanonical genome *k*mer count file size.

#### 3.1.2 Varying the Kmer size with Diverse Database

We assess the performance and run-time of the classifier using 100 reads per genome in a random 5-fold cross-validation experiment against the Standard database.

Supplementary Fig. 2 demonstrates the impact of varying *k*mer sizes on the performance of the “Standard” database across different taxonomic levels. A noticeable trend is the steady increase in recall for higher taxonomic classes, specifically “family” and “genus,” with increasing *k*mer sizes. This suggests that the model becomes more adept at distinguishing between finer, more related taxa as it utilizes longer k-mers. As we show in Supp. Fig. 3, longer *k*mers come at a cost of longer computational time, especially in training but also in testing.

Unexpectedly, the graph indicates a huge drop in performance at higher taxonomic levels, specifically 9-12mer for the superkingdom level. The rate of drop in recall decreases for 14-15mers. The drop in recall rates for the 9-12mers is a point of interest as it suggests a complex interplay between *k*mer size and classification accuracy.

The complex interplay of *k*mers relates to long- and short-range evolution. The database, composed mostly of prokaryotes known for high mutation rates (as compared to eukaryotes), produces results that shows that shorter *k*mers are effective at capturing higher levels of taxonomic classification of the reads, similar to the effectiveness of tetramers in similar scenarios [8]. Longer *k*mers, while capturing specific genomic details (e.g., mutations) of recent evolution, are susceptible to mutations caused by distant evolution, resulting in “class specificity”. Shorter *k*mers are more likely to be less class specific and pick up higher level taxonomic signals. (Please also see Supplementary Appendix 2 for a discussion of the effect of smoothing favoring long genomes).

As indicated in the superkingdom confusion matrix inSupplementary Fig. 5a, we can see that recall rates (for small *k*-mer sizes) are high for Bacteria – however, the other phyla, Eukaryotes/Viruses/Archaea perform worse. At large *k*, recall rates improve for Eukaryotes, but drop for Bacteria. Recall may improve for Eukaryotes because of the fact that Eukaryotes have longer *k*-mers that are more distinctive while prokaryotes have more mutational disruptions. NBC’s imbalance in classification is reflected in the micro vs. macro recall averages (Supplementary Figs. 2 and 4) – since micro-recall calculates on the test data that contains an imbalance of classes (e.g. when simulating from the Standard database, more Prokaryotes are simulated), it can calculate performance based on this imbalance. Therefore, macro-recall is more of a class-balanced measure. Since NBC++ classifies everything, false positives for a predicted class are false negatives for a true class. Therefore, micro-recall and micro-precision are synonymous for NBC++. More detail can be seen in the confusion matrices for common human gut taxa in supplementary Figs. 5a-f.

#### 3.1.3 Kmer sweeps for Prokaryotic Kingdoms

The observations of surprisingly low classification performance (compared to our previous works) led to our hypothesis that the inclusion of the full diversity of life, particularly when considering the marked distinctions between Bacteria and Viruses, might introduce inconsistencies in classification performance. To test this assumption, the experimental procedure was replicated with a modification: the training set was exclusively composed of prokaryotic sequences.

The results when using prokaryotic-only sequences are presented in Supplementary Fig. 6a-c, which reveal an overall positive micro-recall trend across all taxonomic levels in relation to the increasing lengths of k-mers. This observation is similar to previous behavior, albeit at lower performance due to the higher diversity of the database. The superkingdom micro-performance is similar to what we would expect (similar to accuracy rates observed in [11]), and the discerning classifications are further seen in the macro-recall/precision.

#### 3.1.4 Varying the Database Size

We constructed and generated testing reads for the three distinct database configurations: Basic, Standard, and Extended. Subsequent classification tasks were carried out, wherein the testing reads of each database were evaluated against their respective training sets. The results of these experiments are illustrated in Supplementary Fig. 7a-c, which highlight the performance dynamics across these database variations.

A notable observation from the micro-recall/precison graph in Supplementary Fig. 7a is the substantial improvement in classification performance when transitioning from the Basic to the Standard database configuration. While using the Standard database leads to the best micro-recall, this result is due to the heavy bias in prokaryotes in the test dataset, which the classifier trained with the Standard database is better at classifying. The macro-recall graph (Supp. Fig. 7b) show that recall improves when using Standard database (compared to the Basic database). Furthermore, we observe very little improvement in macro-precision (Supp. Fig. 7c) with the Standard database, which itself improved with the use of the Extended database. These observations show that NBC++ performs better at superkingdom level, but its performance drops off for finer taxonomic levels.

#### 3.1.5 Query input data size vs. time

To validate the linear computational time of NBC++, we conducted an experiment to monitor the runtime of the algorithm while increasing the amount of data. This experiment fixed the k-mer size at 9 and utilized the Standard database for classification. The computational resources allocated for this experiment included a fixed system with 180 GB of RAM and 48 processing cores allocated for this. The results, depicted in Supp. Fig. 8, affirm that the classification time scales linearly with the size of the input file.

Further investigation was undertaken to evaluate the classification task on a typical personal computer configuration, characterized by 64 GB of memory and 8 processing cores. An input query file of 240 MB was selected for this purpose, and its classification, using 9mers against Standard database, took 47.3 hours, approximately 6.3 × longer than 180GB/48-core runs that we did, again showing that the main factor affecting wall-clock time is the linear relation to multi-threading linearly, and that there is a slight overhead due to memory partitioning.

### 3.2 Kraken2 Results for Comparison to NBC++

For all datasets, genomes were first added to a Kraken2 library sequentially (to decrease the wallclock time of this step, the genomes in extended folds 2-5 were added through five parallel processes; see Kraken2 add-to-lib in the Appendix for more details). Then, a custom Kraken2 database was built using default settings for each fold. To minimize the number of unclassified reads in testing, the confidence parameter (`--confidence`) was set to 0.001, a low enough value to result in classifications with just a single *k*mer match to a reference sequence.

Given that NBC++ performance was significantly lower for a diverse database of Prokaryotes, Eukaryotes, and Viruses, we wanted to evaluate how another classifier, Kraken2, would perform with the same training databases (see Supp. Figs. 9a-d). It is important to note with Kraken2 that the micro-precision measure now becomes different than micro-recall, since Kraken2 filters out a lot of the reads as “unclassified”. In the micro-recall/precision analysis, unclassified reads are considered as false negatives and are never considered as false positives (and are therefore micro-precision diverges from micro-recall). Kraken2 achieve high precision performance, however, its recall performance is poor. On the other hand, Kraken2 struggles with a diverse database, and only achieves a little over 30% in macro-recall for the Extended database and 10% or less with the Basic database (with better precision and similar recall performance to that of NBC++. Kraken2 vs. NBC comparison results can be found in Supp. Figs. 10a-b, where we observe that Kraken2 exhibits superior performance in general for macro-measures (except for the superkingdom level for Prokaryotes), particularly as the database size expands.

Regarding time efficiency of both classifiers, supp. Fig. 11 provides an overview of the CPU time for a complete run, encompassing both the training and classification phases. We observe that NBC++, when employing a 9-mer approach, demands significantly less training time compared to Kraken2. However, the majority of Kraken2’s time expenditure is allocated to the database construction phase. Once the database is established, Kraken2 demonstrates remarkable speed in the classification of new input queries, outpacing NBC in this respect. This delineation suggests that while NBC is efficient in a holistic sense, Kraken2’s architecture allows it to excel in rapid query classification following the initial database setup.

### 3.3 Human Gut Sample Classification vs. Database Depth

To analyze a human gut sample, we used *k*=9, and examined the results across the three database depths (Basic, Standard, and Extended), shown in Fig. 1 and supp. Fig. 12. Fig. 1 also shows the Bray-Curtis dissimilarity metric to gauge the comparative similarity between the classified compositions of the human gut sample across the different databases. We see that Standard and Extended databases have more concordance than Basic and Standard databases, which is expected due to the sparse representation of the Basic database. We note that the Standard database seems to lack good representation of viruses, as shown for the superkingdom analysis (Fig. 12a). Uroviricota are found in significant quantities in both Basic and Extended databases, with Basic indicating 3% more RA than Extended. Because of few viral labels at the order level, Standard and Extended databases are very similar.

**Figure 1:**
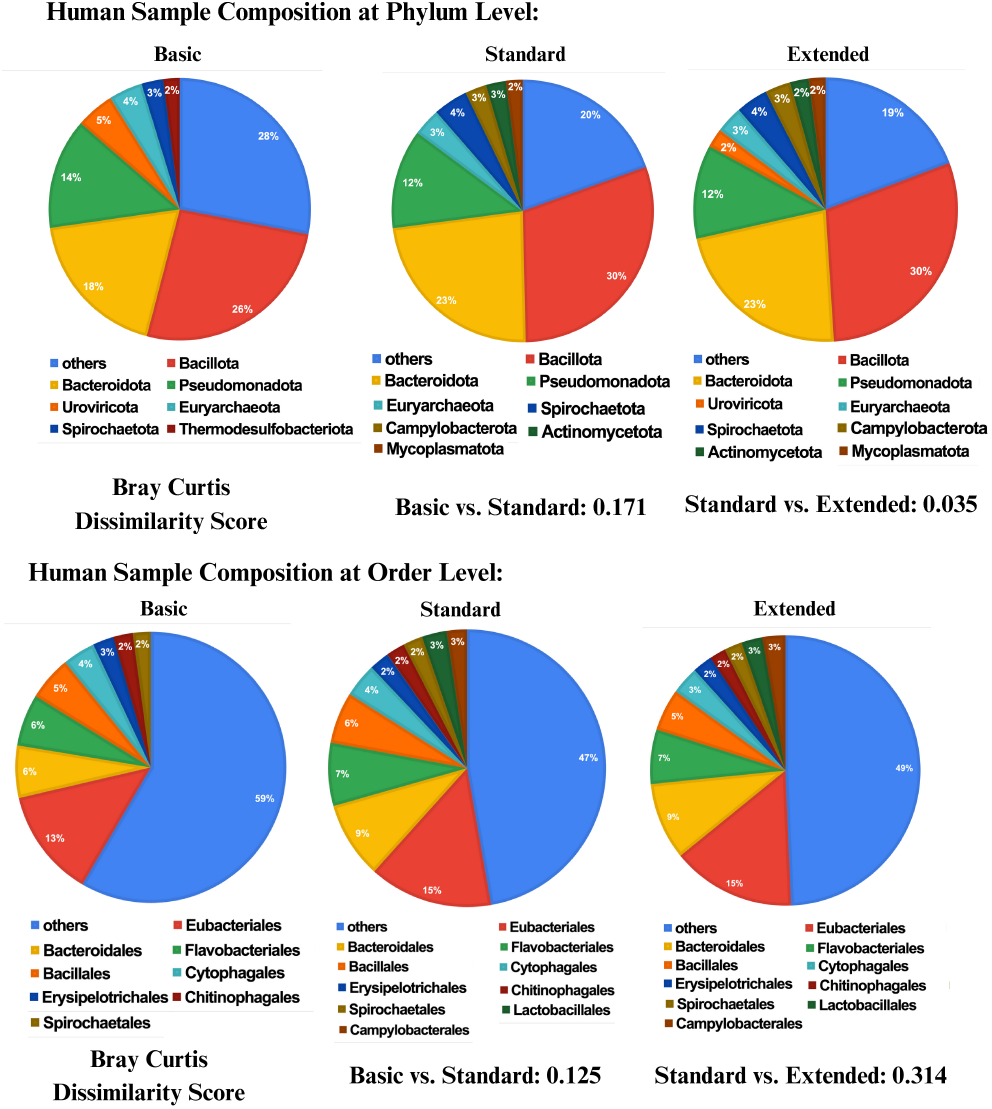
The Phylum and Order composition of the sample with taxa over 2% relative abundance shown. Standard and Extended have more concordance than Basic and Standard.

## 4 Conclusion

We show that NBC++ can now be queried with realistic, large datasets. NBC can provide reasonable classification with a minimal database, especially in identifying particular superkingdoms and phyla, highlighting its ability to differentiate classes using *K*mer frequency. NBC++ may have less training time than some classifiers but at a cost of more testing (inference) time.

We tested a diverse database of at least one example per genera (basic), one (or a few) per species (Standard), and one with more depth of examples (Extended). While it was thought that such a database would represent the tree of life, we found that even a state-of-the-art classifier struggled with training on prokaryotes, viruses, and eukaryotes at the species level. Hopefully the study within the paper provides a jumping point for further studies to examine the effect of breadth and depth of databases on taxonomic classifiers.

We should also note that an important improvement is the ability to obtain all log-likelihoods (of each read against each database genome). This ability enables future work of implementing a confidence measure and/or novelty detection using the log-likelihood scores determined by NBC++.

## Supporting information

All supplementary figure files

The assembly summary files for the depth databases used in the experiments

## 6 Author contributions statement

H.D. and G.L.R. conceived the experiment(s), H.D. conducted the NBC experiment(s), G.H. conducted the Kraken2 experiments, H.D., G.H., and G.L.R. analysed the results. All authors wrote and reviewed the manuscript.

## 7 Acknowledgements

We are thankful for computational time from the Drexel University Research Computing Facility. We would like to thank M. Saleh Refahi for providing insights during the development process.

## 9 Data availability

NBC++ source code is available at http://github.com/EESI/Naive_Bayes.

A docker container is available at: https://hub.docker.com/r/eesilab/nbc_complete_toolset. The classifier training data trained on Basic/Standard/ Extended databases using canonical 9-mers and the associated assembly summary files of the databases are available at https://zenodo.org/records/11657719.

Summarized data results are at: https://zenodo.org/uploads/11643985. The human sample reads are located at SRA ID: SRS105153.

## Appendix

The buffer sizes are calculated using the following formulas:

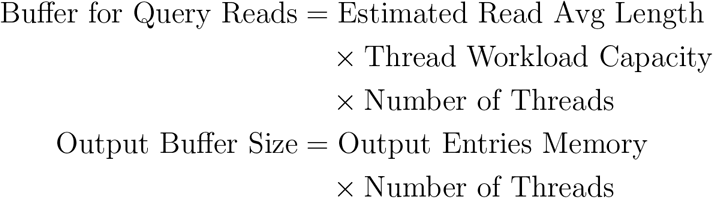

### 9.0.1 Trained Class Memory Allocation

Memory for loading classes is dynamically allocated based on the total memory specified by the user, divided by the average memory requirement for each class derived from training files, plus any additional temporary memory needed during processing. This temporary memory requirement is approximated as follows:

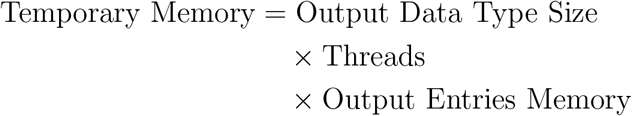

#### 9.1 *K*mer Smoothing may favor longer genomes that tend to have more diverse *k*mer vocabularies by chance

Reconsidering the analysis from the preceding section, the natural logarithm of the vocabulary size for different k-mer lengths can be calculated as follows: for 9-mers, the upper limit to the Vocabulary value is computed using the expression ln 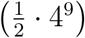, yielding approximately 11.784. For 13-mers and 15-mers, the respective expressions ln 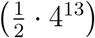 and ln 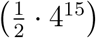 result in values of 17.329 and 20.101, respectively. This delineates a trend wherein the incremental differences between the logarithmic values diminish as the k-mer length increases.

Adapting the Naive Bayes’ theorem into a rearranged form yields the equation:

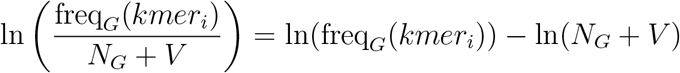

Within this context, for genomes of considerable length, the propensity to include specific k-mers found within the reads escalates. As the k-mer length extends, the natural logarithm of the vocabulary size, a pivotal factor in the subtractive segment of the equation, starts to converge with this subtractive element, thereby favoring longer genomes with elevated scores.

### 9.1.1 Kraken2 add-to-lib

Because all training dataset genomes were stored in separate files, a bash script was used to sequentially apply the add-to-library function from kraken2. This took a reasonable amount of time for the basic/standard datasets, but took multiple days when applied to the extended dataset. To get around this issue, the dataset was divided into 5 equivalent chunks. This allowed each chunk to be added simultaneously, reducing the wallclock runtime to 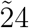 hours from 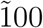 hours. The kraken2 was then built on each library using default settings - the only exception being the extended database, which was run using 48 cores (all other jobs were done with a single core) to reduce the runtime. The taxonomy file was downloaded early on in testing, and subsequently cloned for each database.

#### Assembly Summary Files for the Basic, Standard, and Extended Databases

The files are in the database assembly_summaries.zip, with the format of {database}_assembly_summary.csv.

