## Supplementary material for "The Naïve Bayes Classifier++ for Metagenomic Taxonomic Classification – Query Evaluation": All supplementary figure files

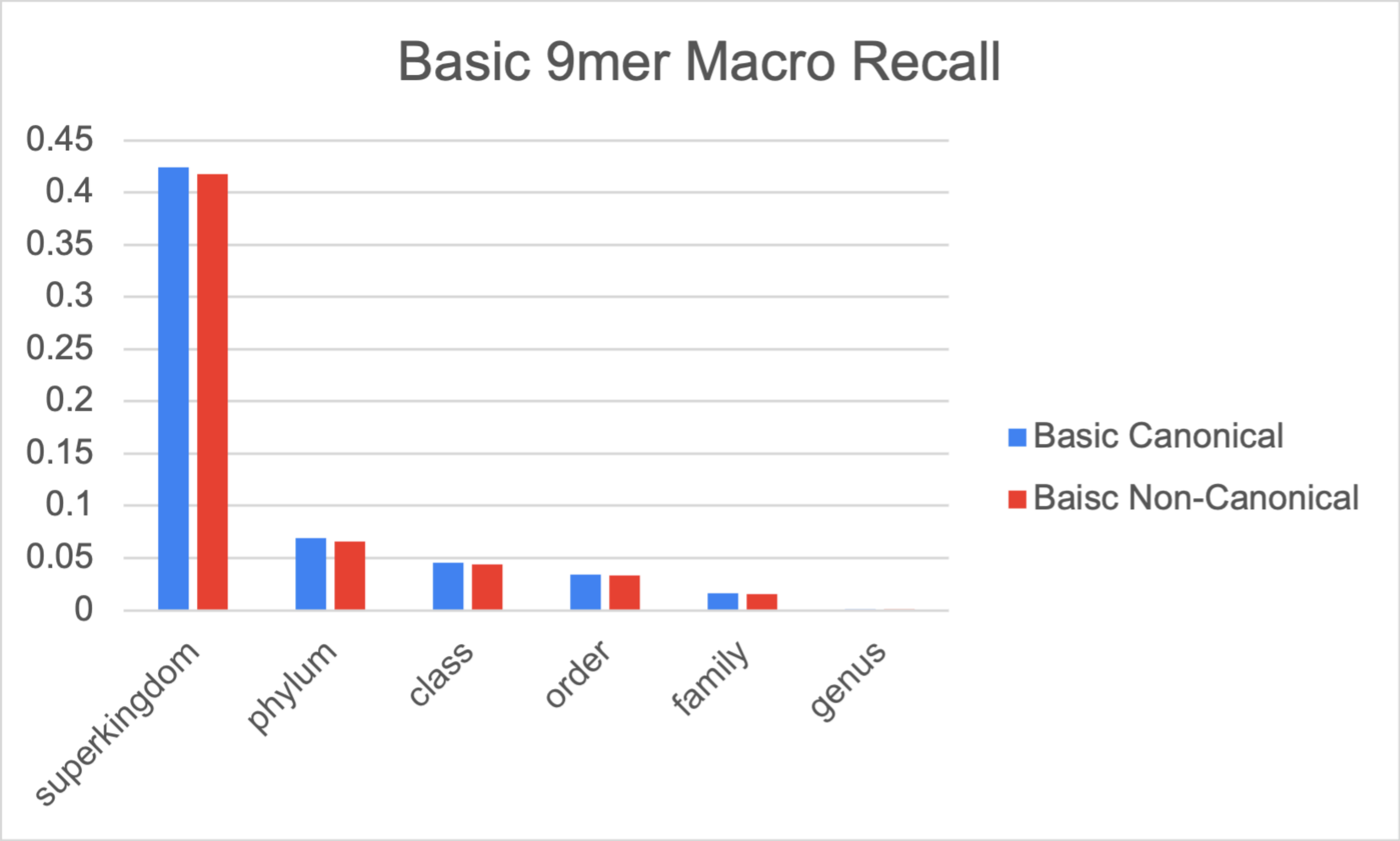


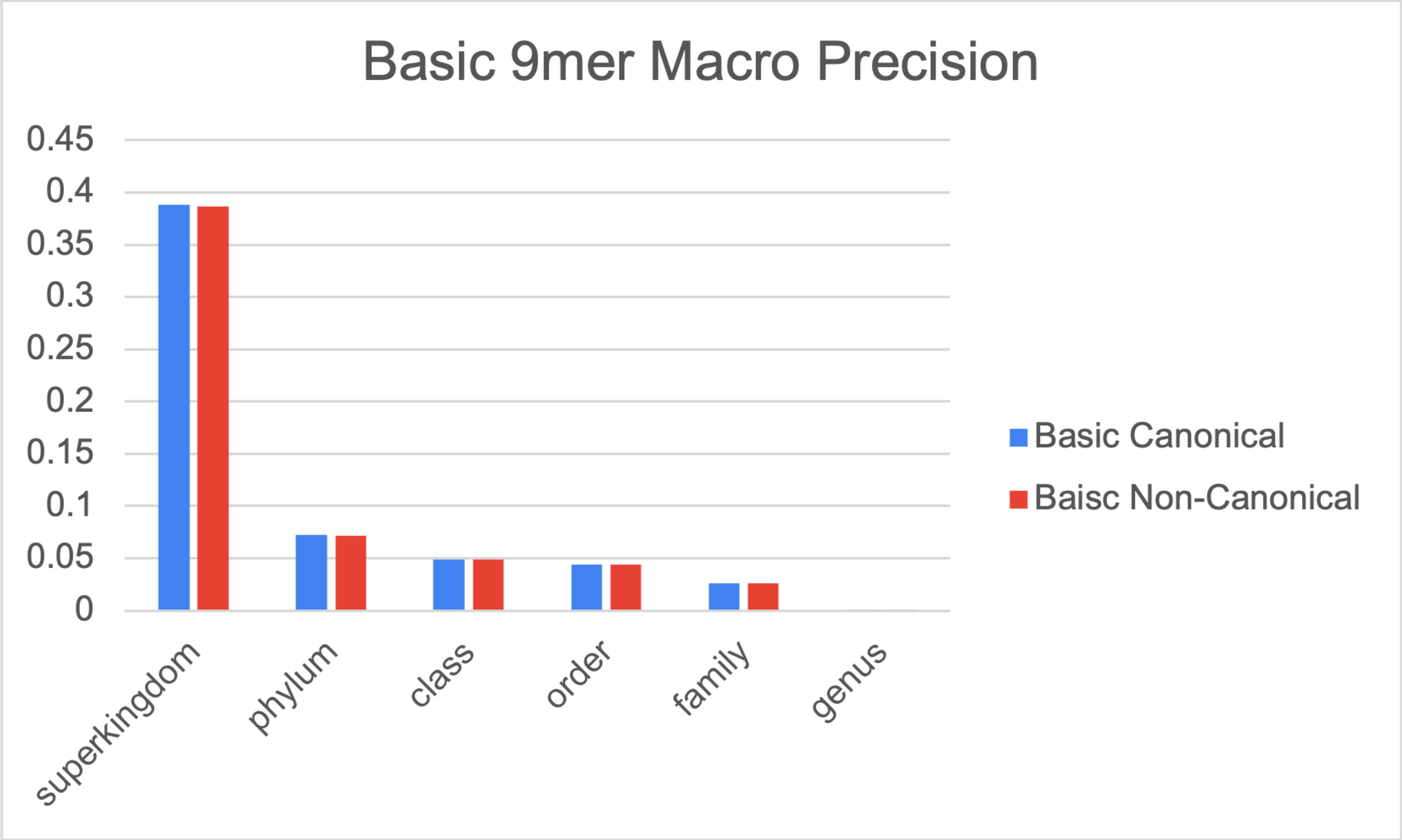


**Fig. 1: Macro-Precision and Recall for Canonical vs. Non-canonical vs. taxonomic level for the Basic Database at 9mers. There is little difference in using different canonicality.**

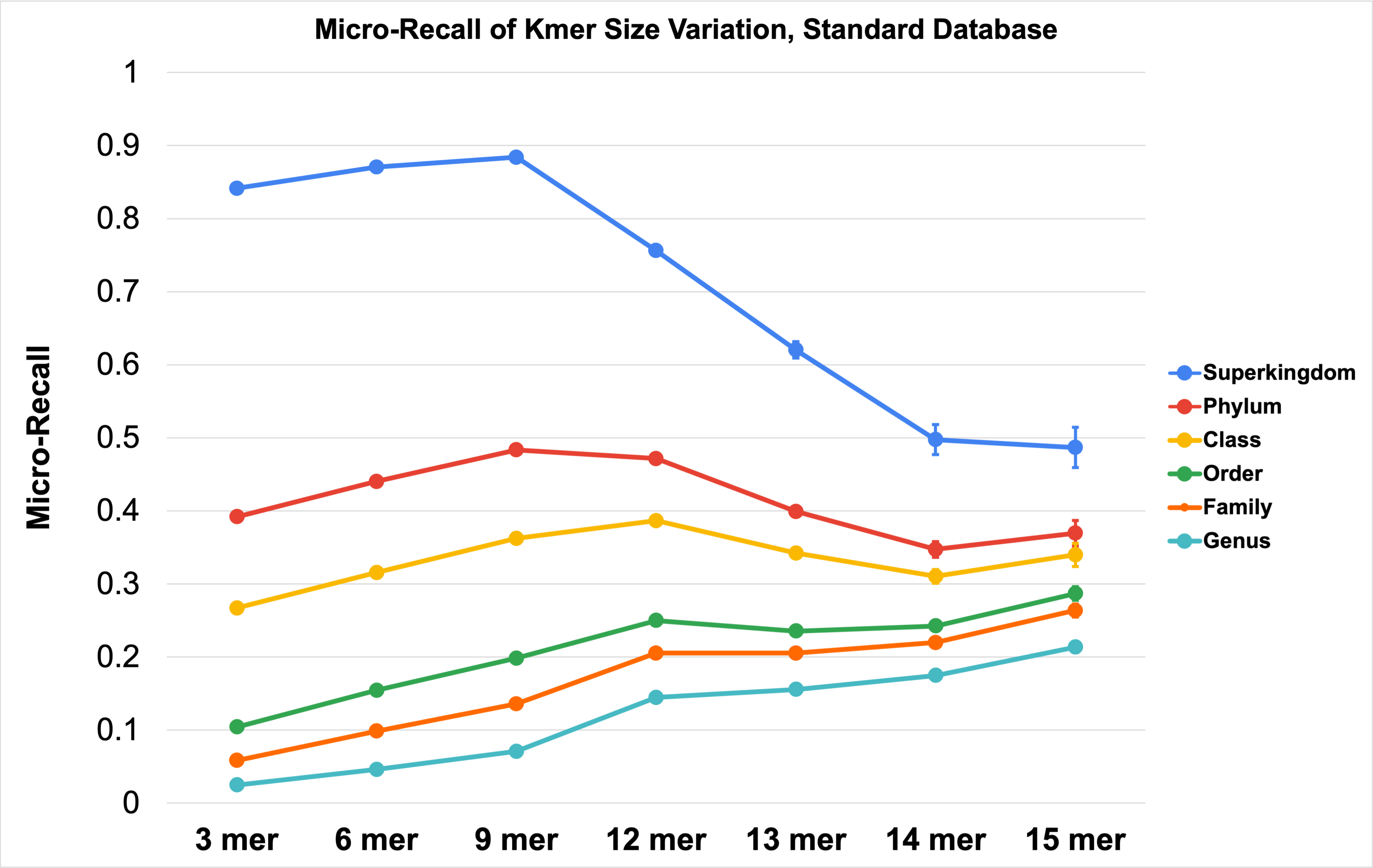


**Fig. 2: Micro-Recall (Micro-Precision) vs. NBC Kmer Size for the Standard database.**

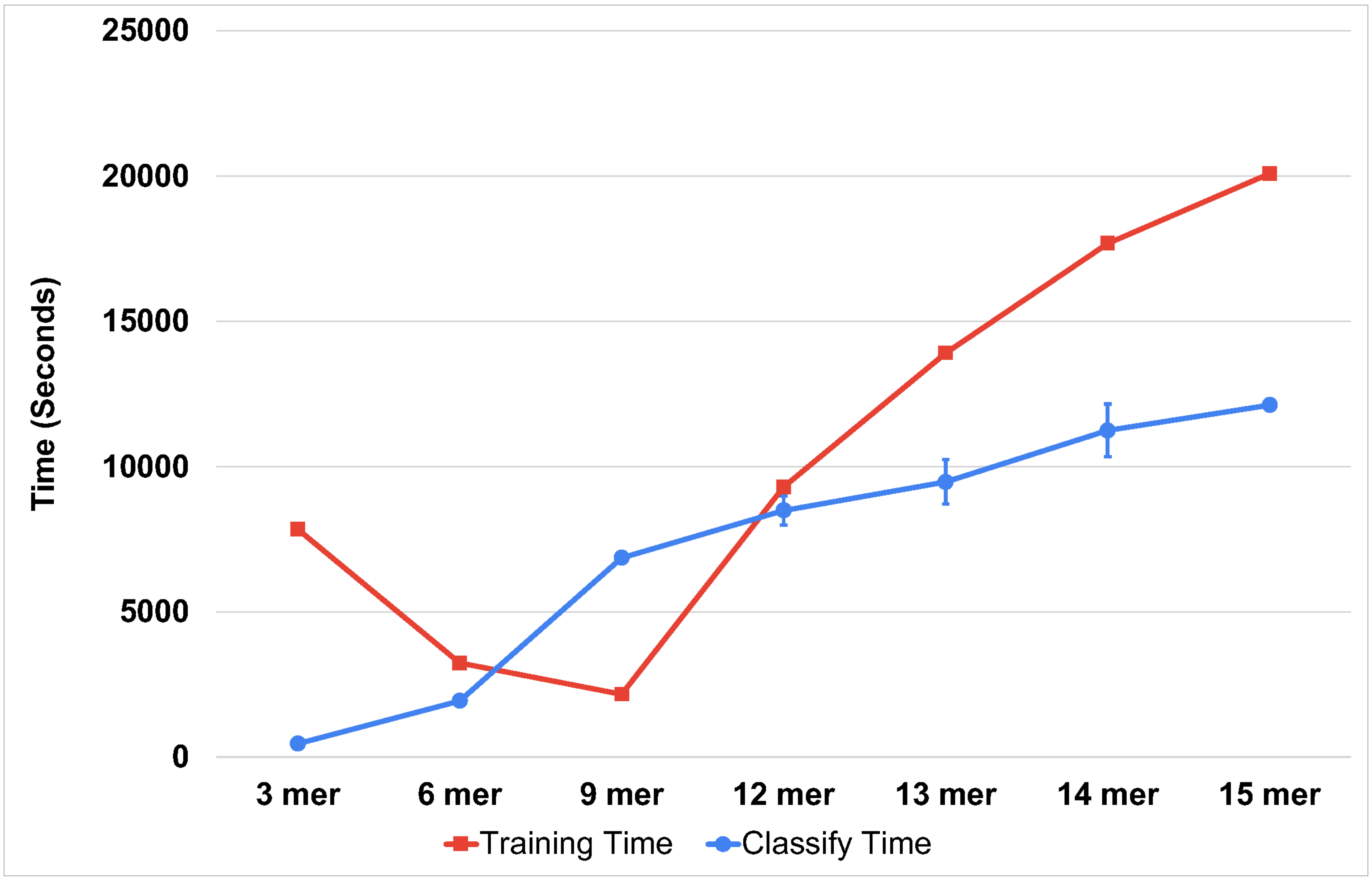


**Fig. 3: Training/Classify Time vs. NBC Kmer Size for the Standard database**

**Average Time in Seconds**

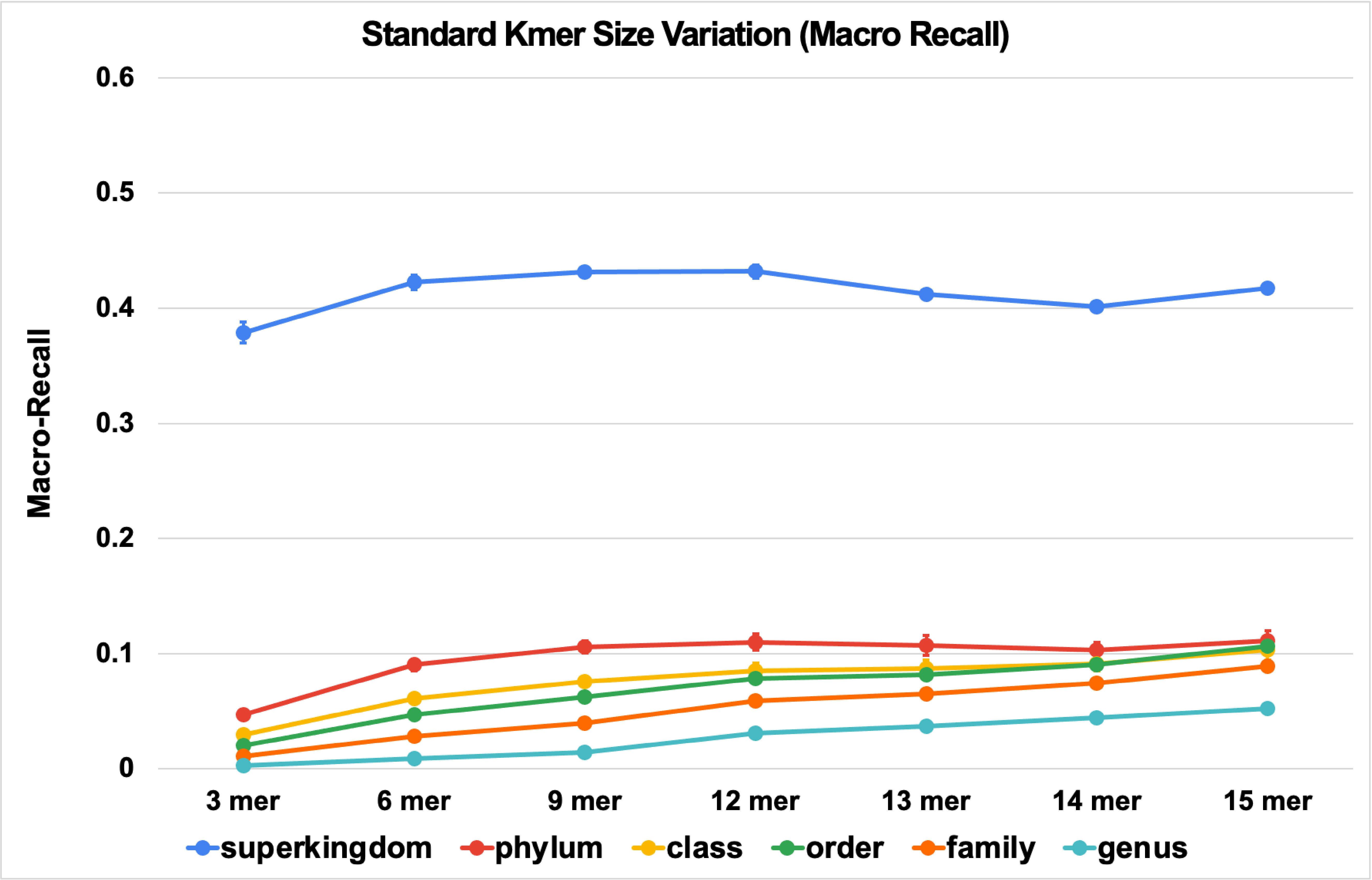


(a)

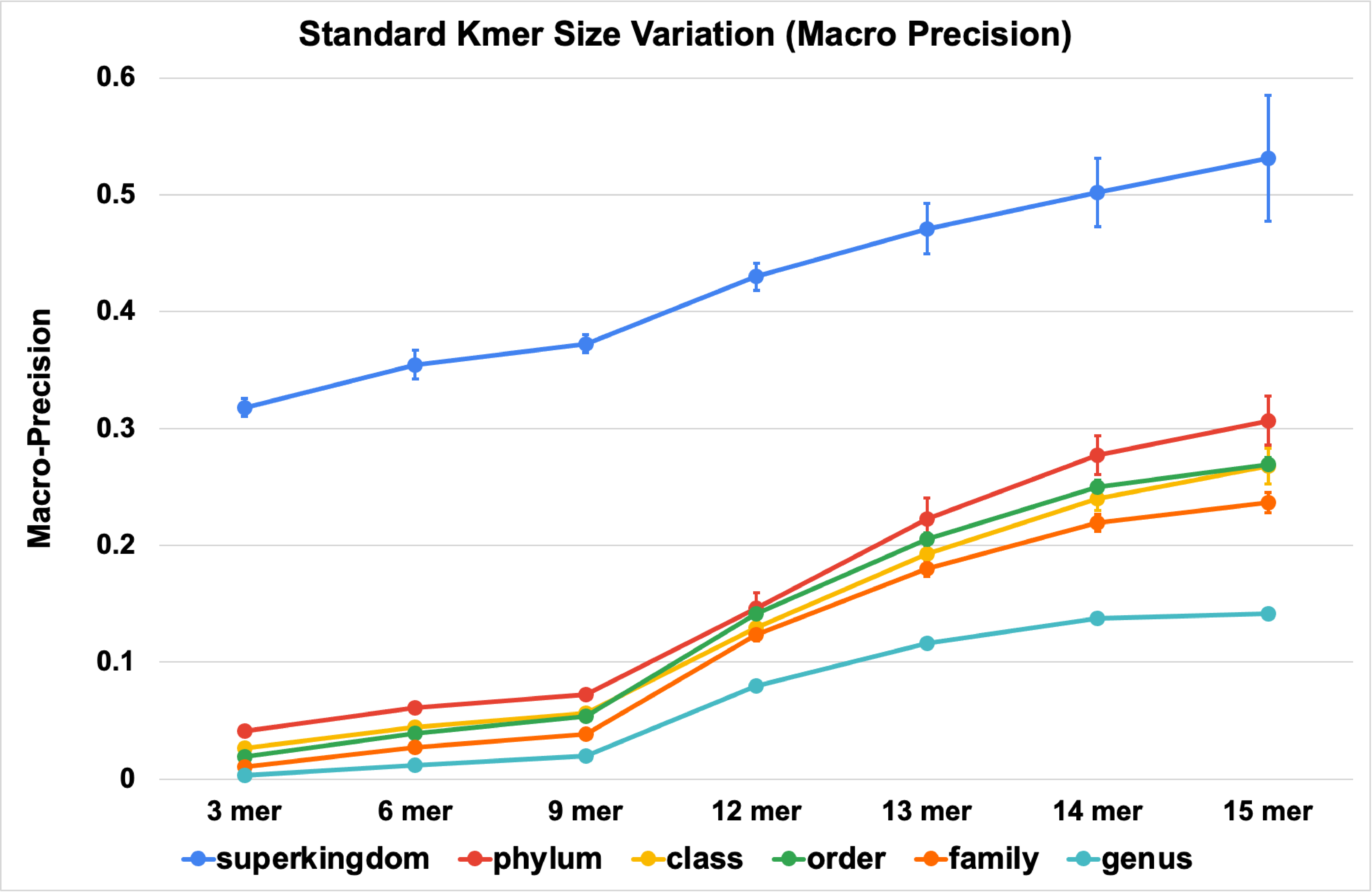


(b)

**Fig. 4: Macro-Recall/Sensitivity (a) and Macro-Precision (b) vs. NBC Kmer Size for the Standard database.**

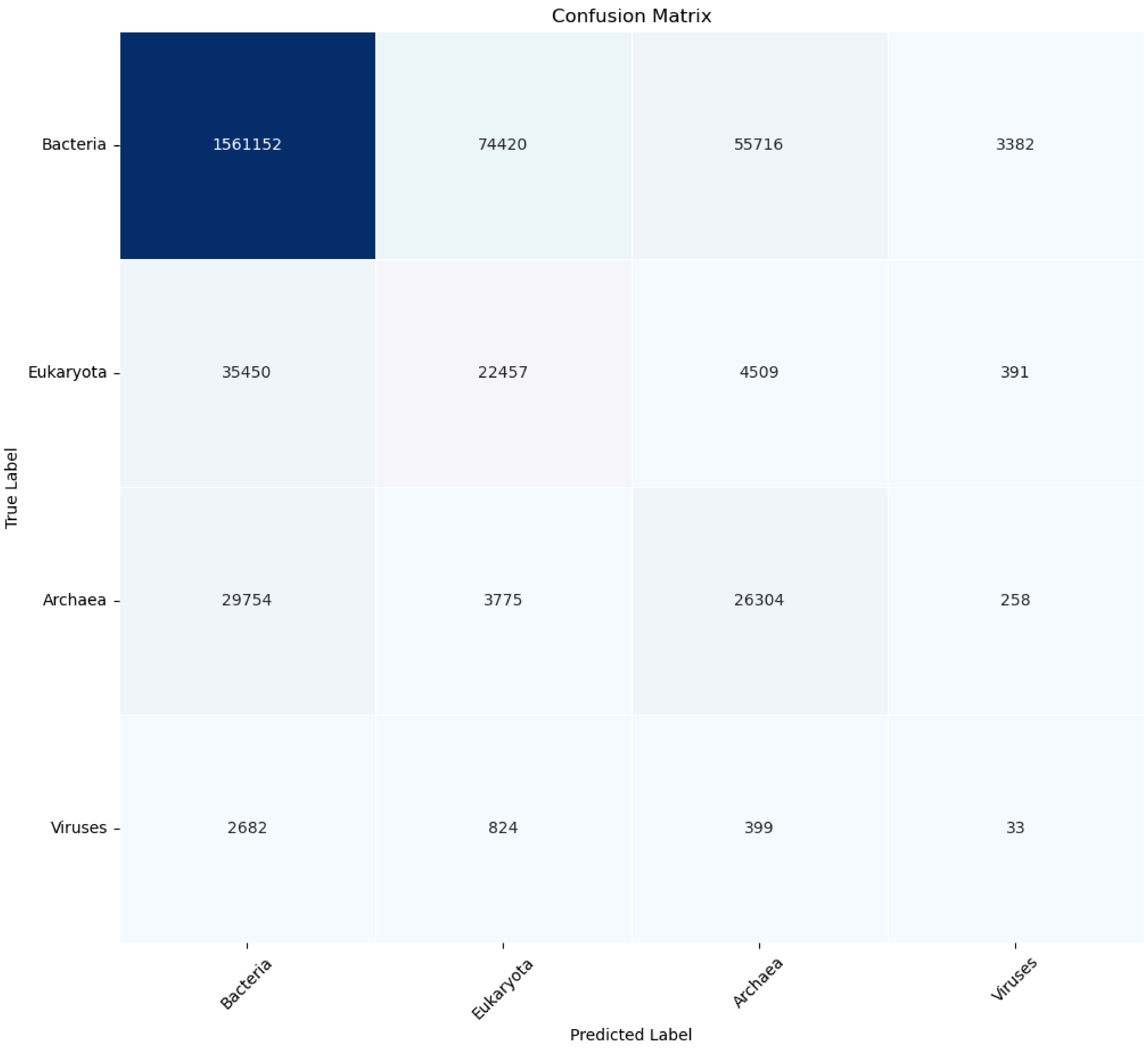


**Fig. 5a: Confusion Matrix of NBC 9mer, Standard Database Superkingdom Level**

**Values are taken from the average of the 5 folds.**

**Only taxa with high concentrations in the human sample result are selected for illustration**

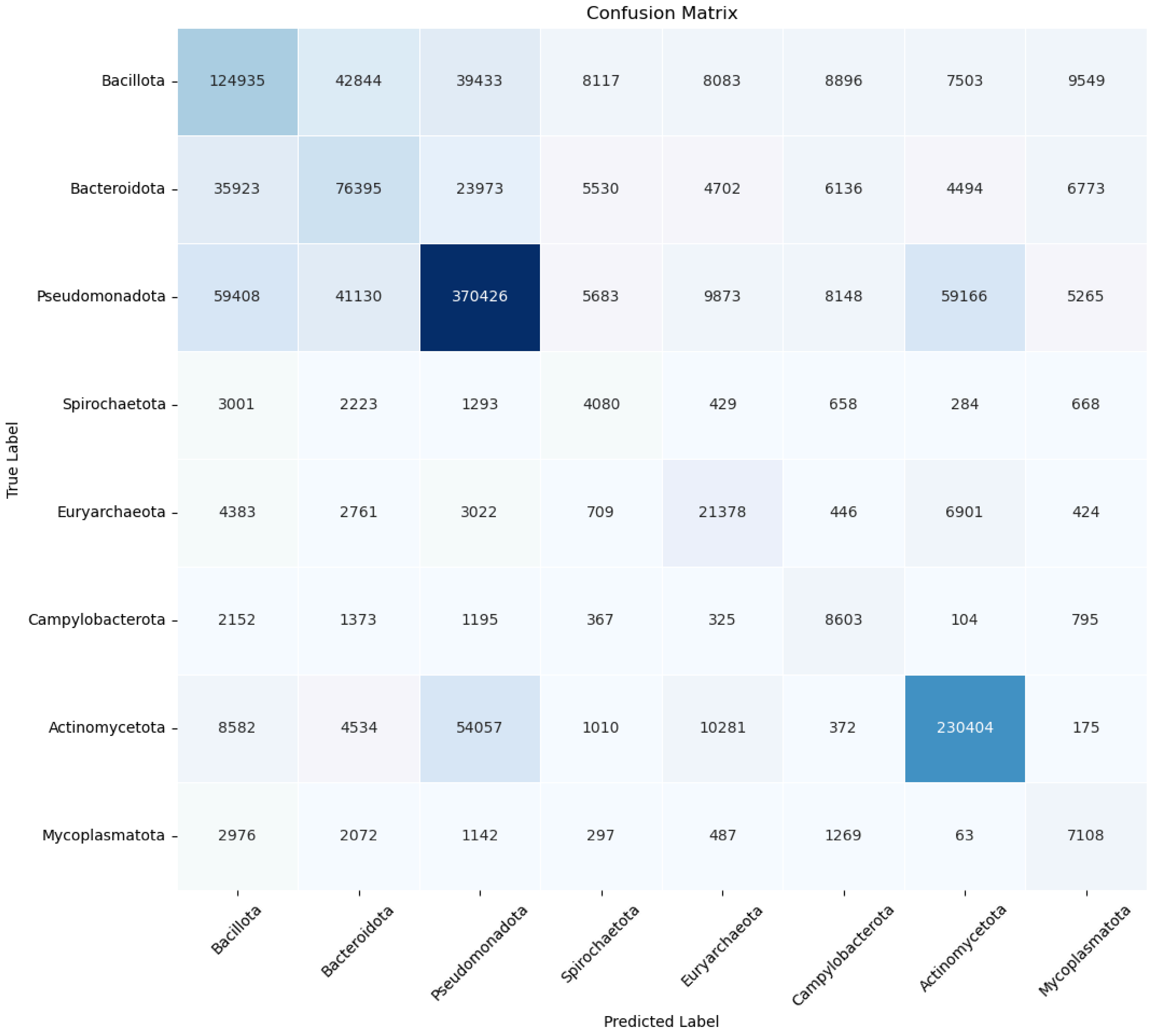


**Fig. 5b: Confusion Matrix of NBC 9mer, Standard Database. Phylum Level**

**Values are taken from the average of the 5 folds.**

**Only taxa with high concentrations in the human sample result are selected for illustration**

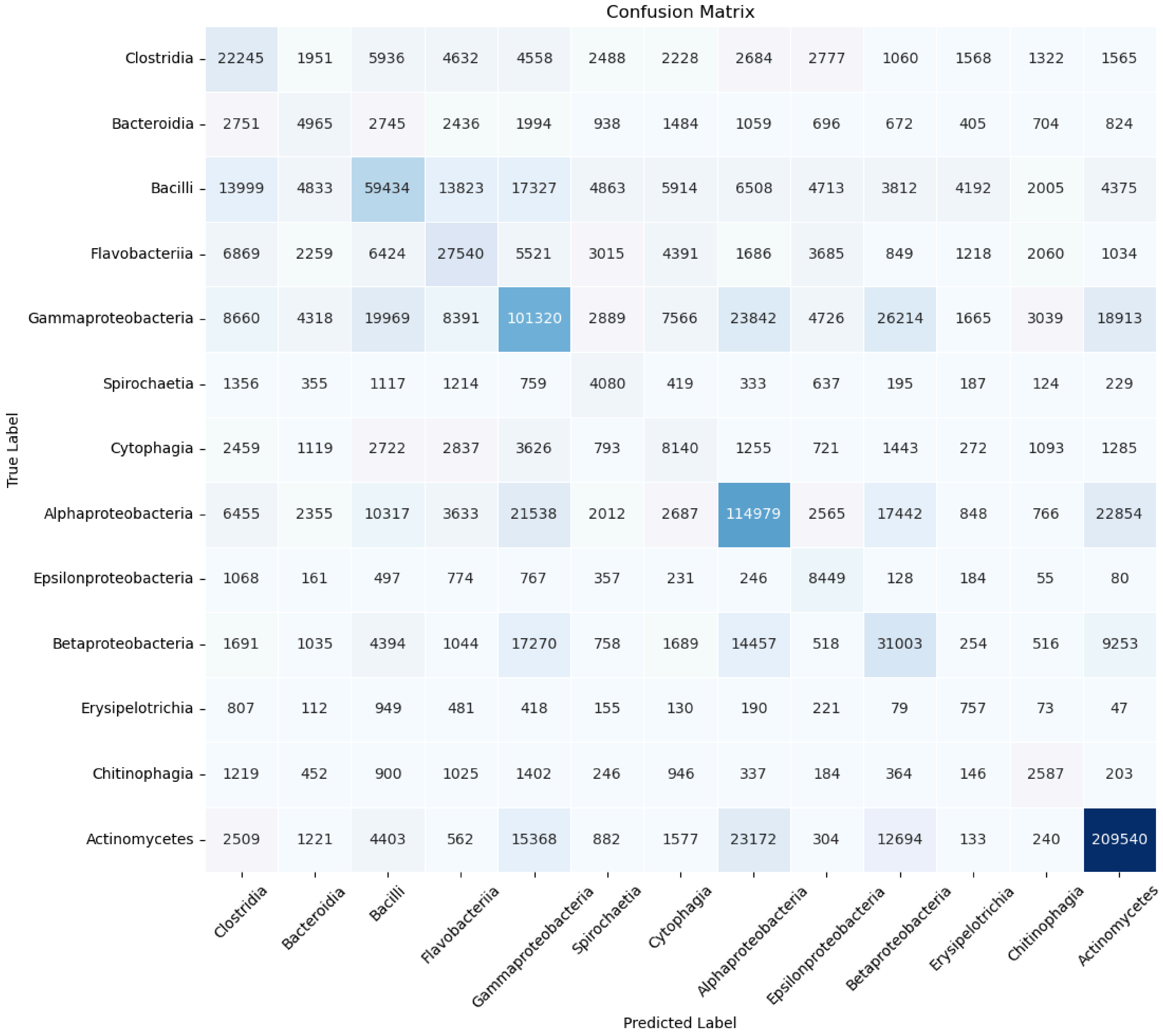


**Fig. 5c: Confusion Matrix of Standard, 9 mer.**
**Class Level**
**Values are taken from the average of the 5 folds.**
**Only taxa with high concentration in the human sample result are selected for illustration**

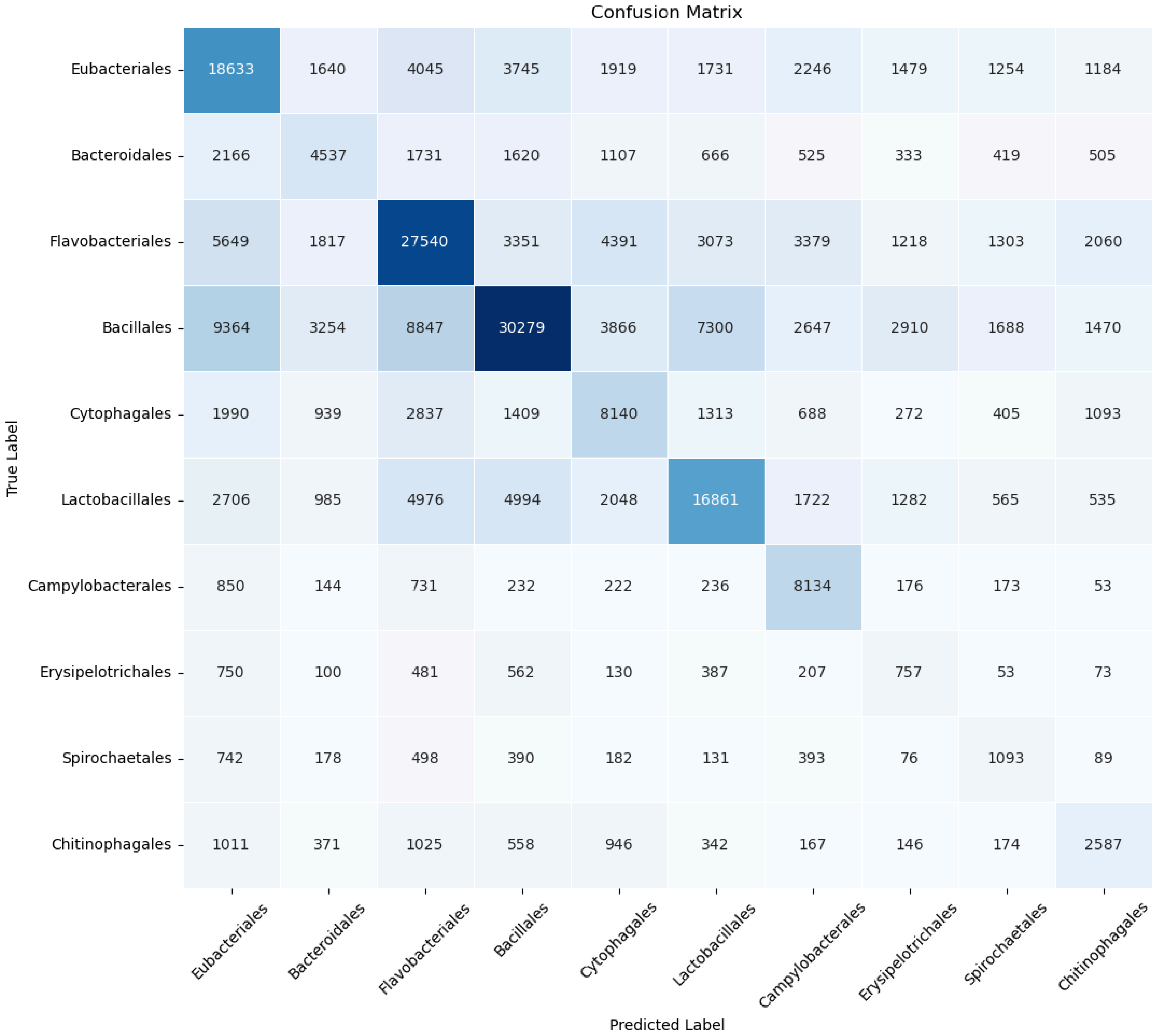


**Fig. 5d: Confusion Matrix of Standard, 9 mer.**
**Order Level**
**Values are taken from the average of the 5 folds.**
**Only taxa with high concentration in the human sample result are selected for illustration**

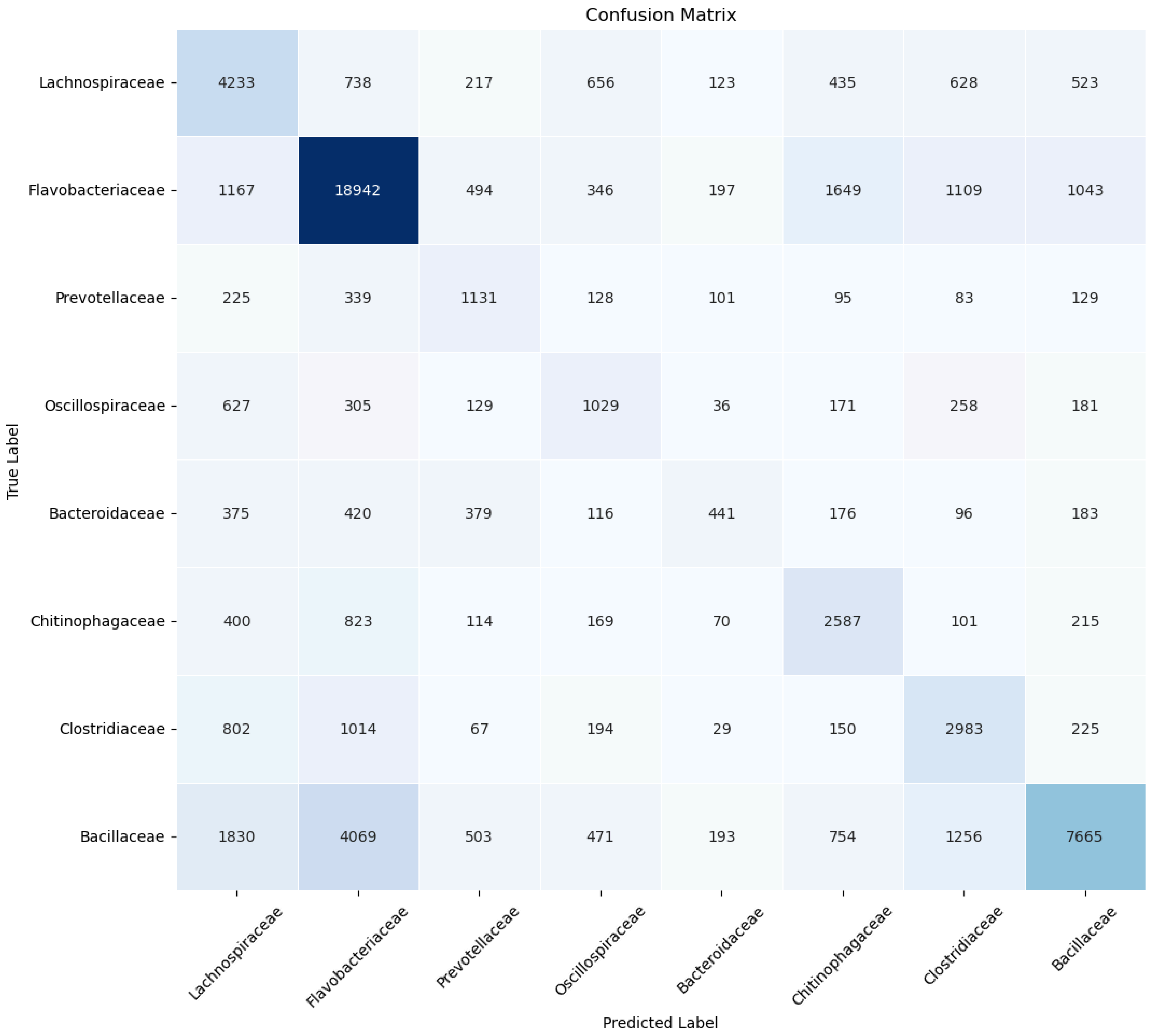


**Fig. 5e: Confusion Matrix of Standard, 9 mer. Family Level**
**Values are taken from the average of the 5 folds.**
**Only taxa with high concentration in the human sample result are selected for illustration.**

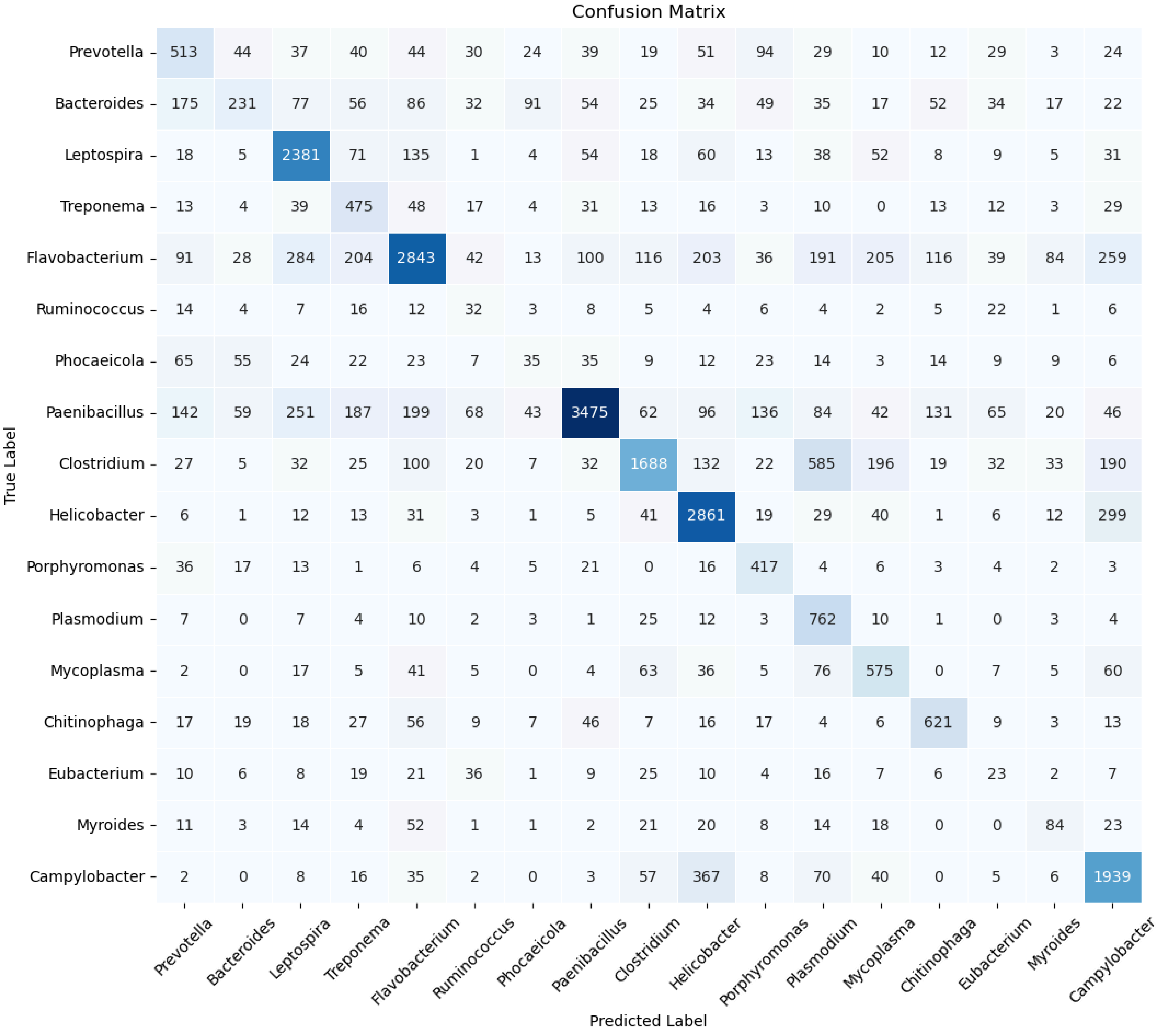


**Fig. 5f: Confusion Matrix of Standard, 9 mer Genus Level**
**Values are taken from the average of the 5 folds.**
**Only the group with high concentration in the human sample result are selected**

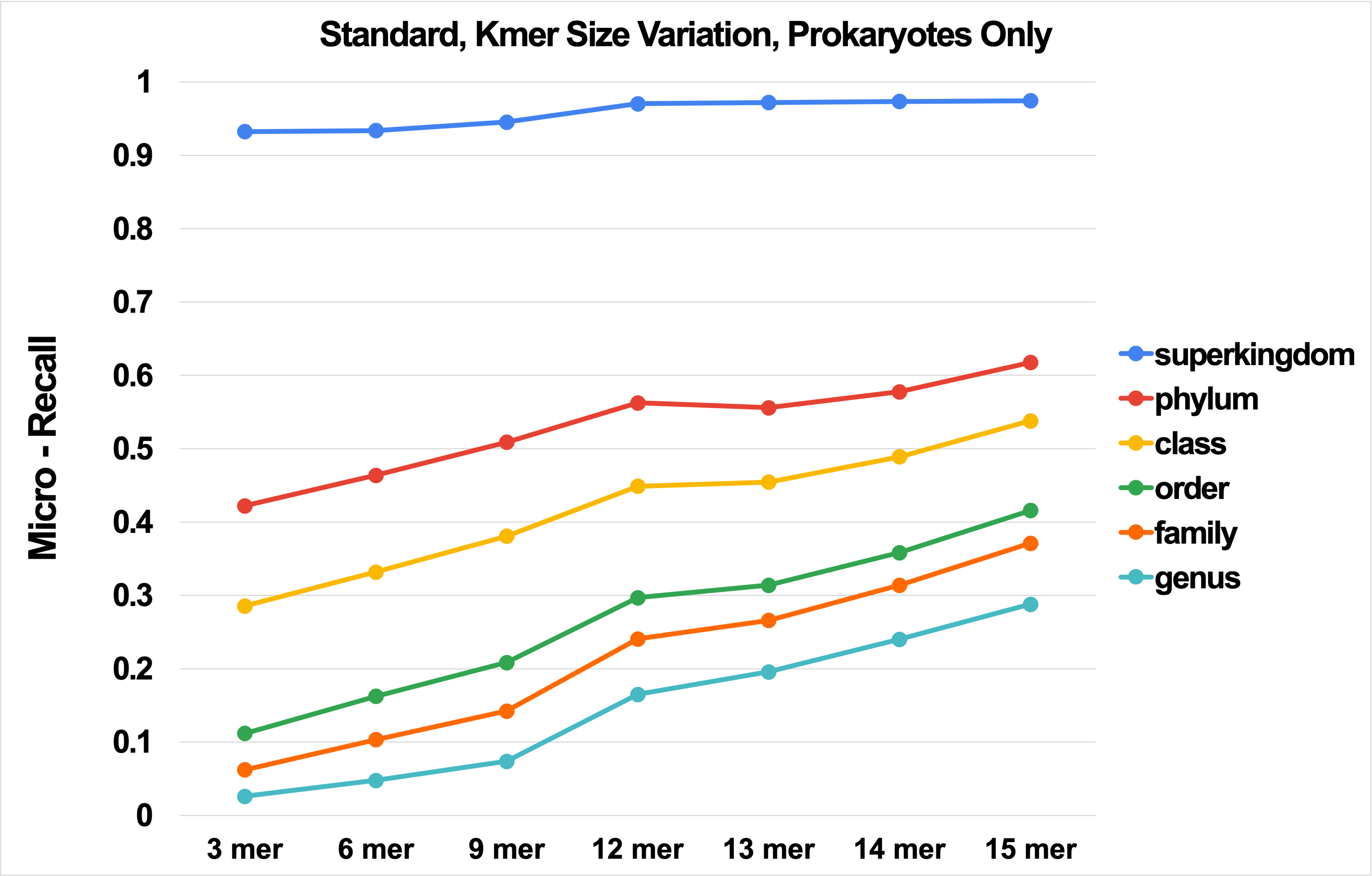


**Fig. 6a: Micro-Recall (Micro-Precision) vs. NBC Kmer Size for the Standard database, Prokaryotes Only.**

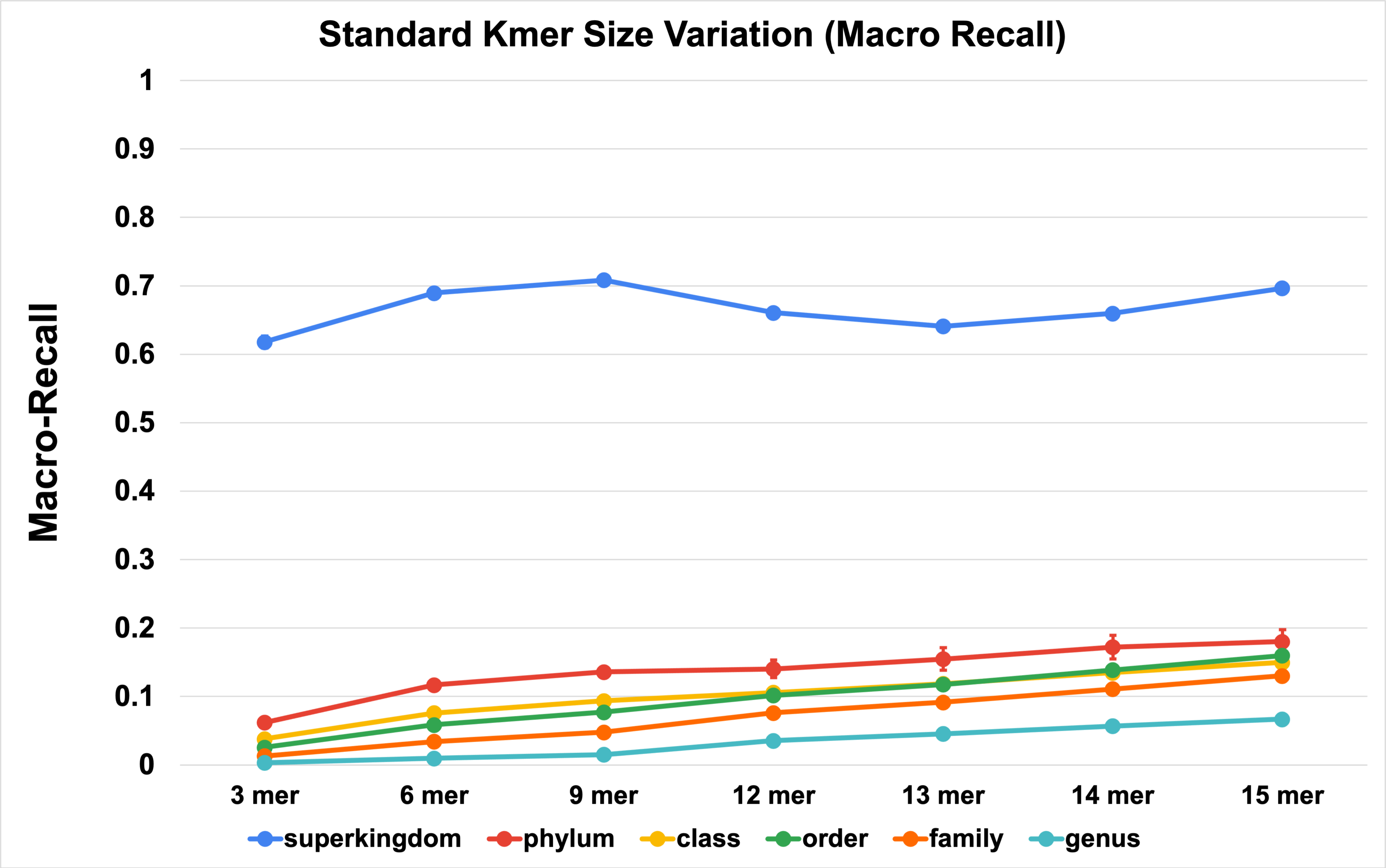


**(b)**

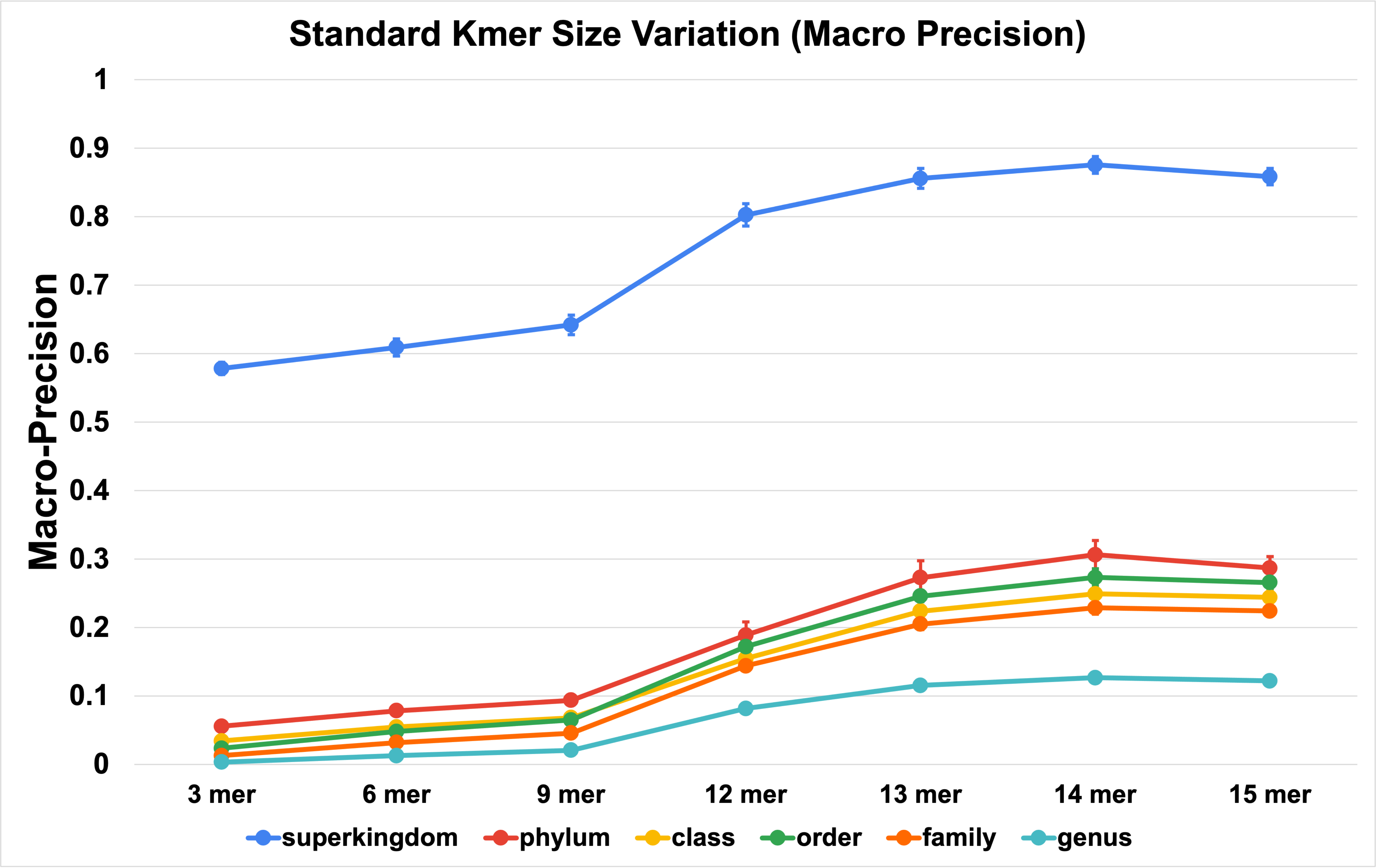


**(c)**

**Fig. 6: Macro-Recall/Sensitivity (b) and Macro-Precision (c) vs. NBC Kmer Size for the Standard database, Prokaryotes Only**

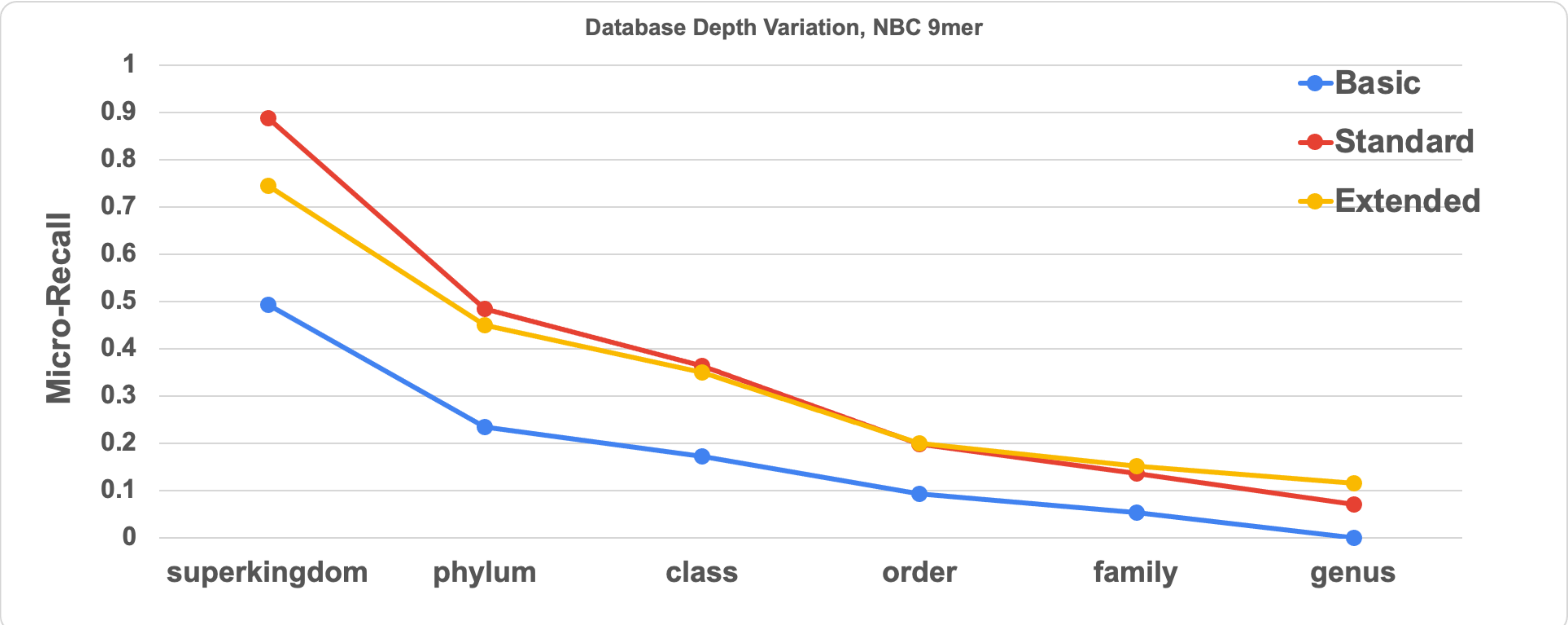


**Fig. 7a: NBC-9mer Micro-Recall/Sensitivity (Micro-Precision) vs. Taxonomic Level vs. Database Depth**

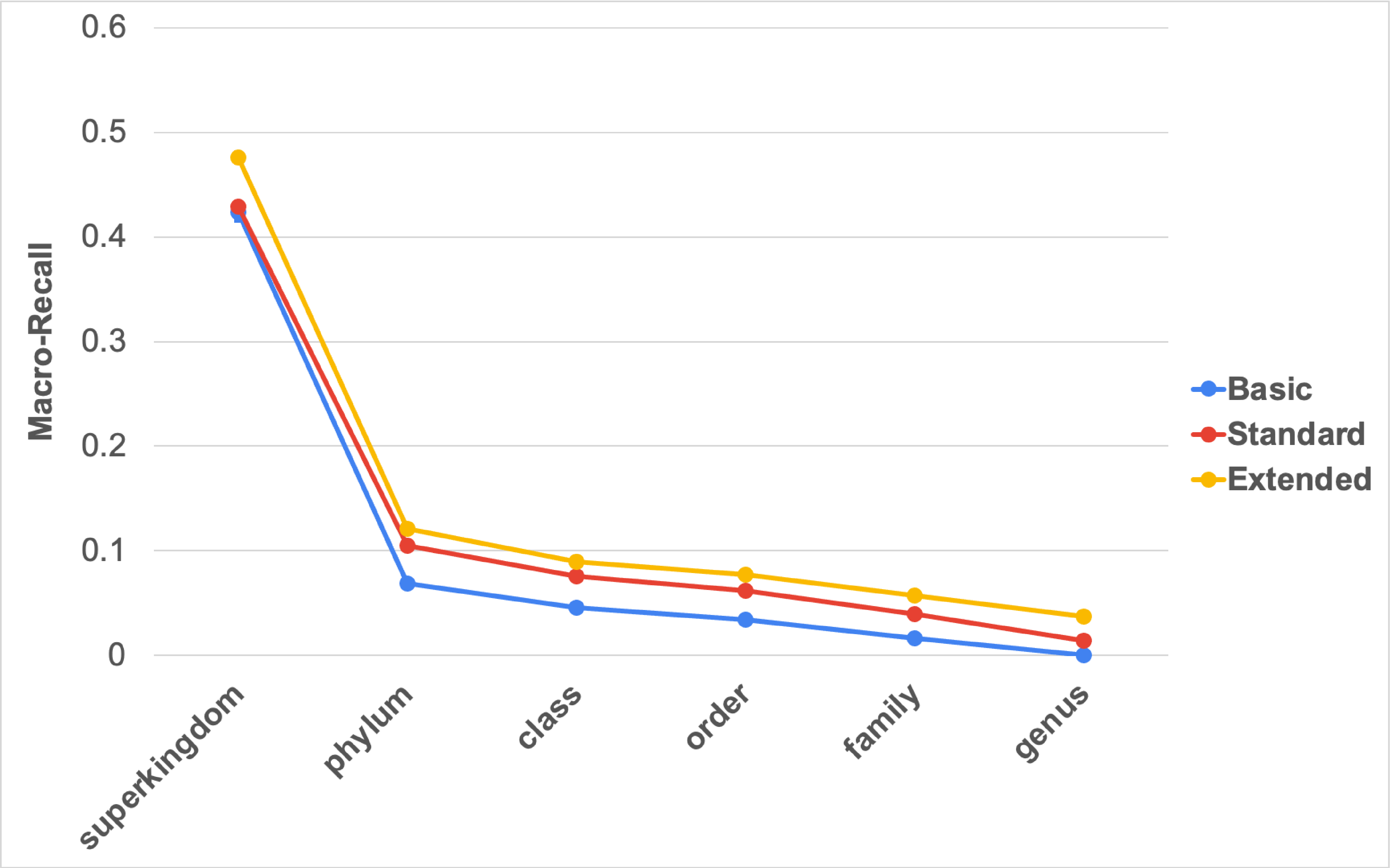


**(b)**

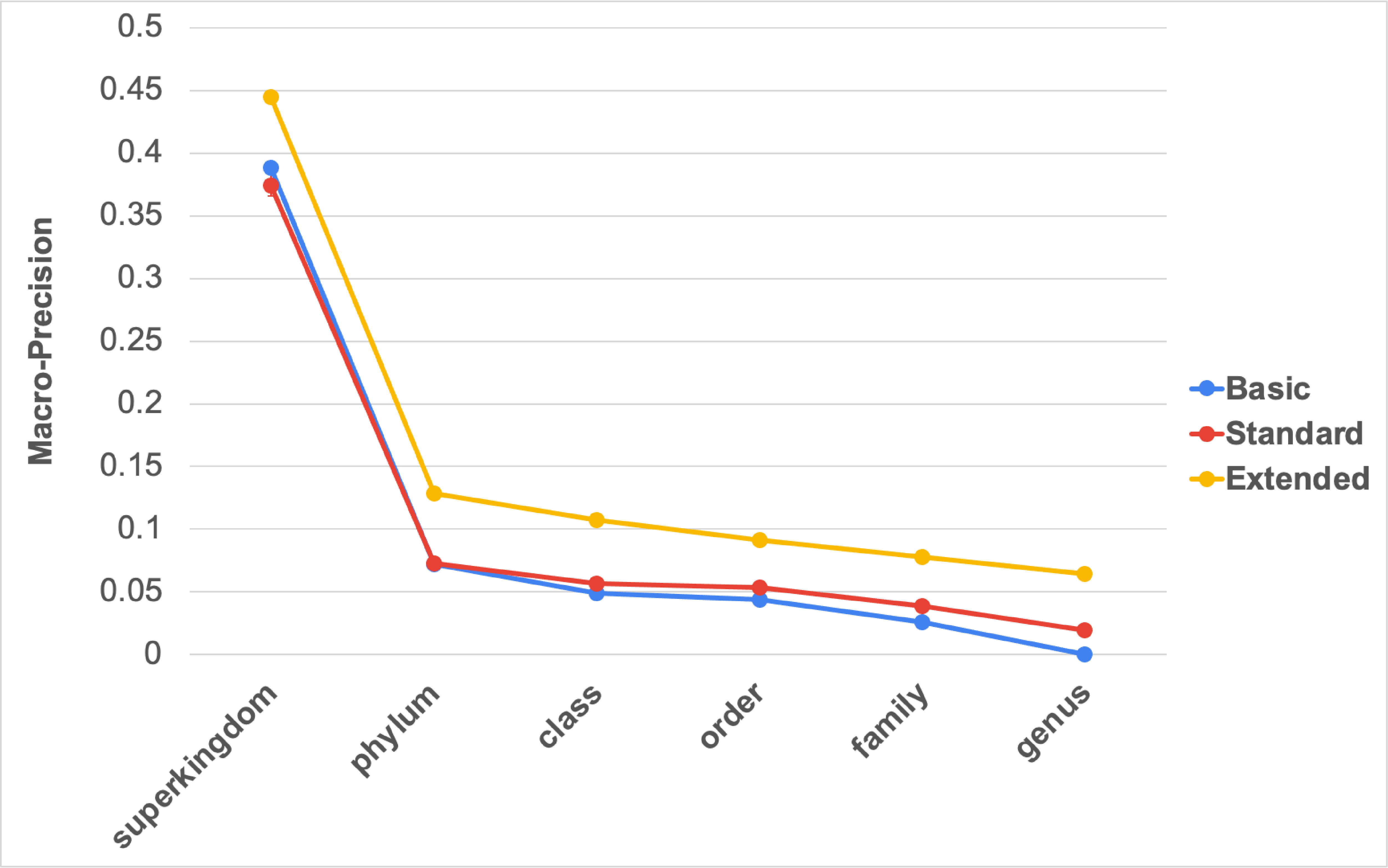


**(c)**

**Fig. 7: NBC (for 9mers): Macro-Recall/Sensitivity (a) and Macro-Precision (b) vs. Database Depth**

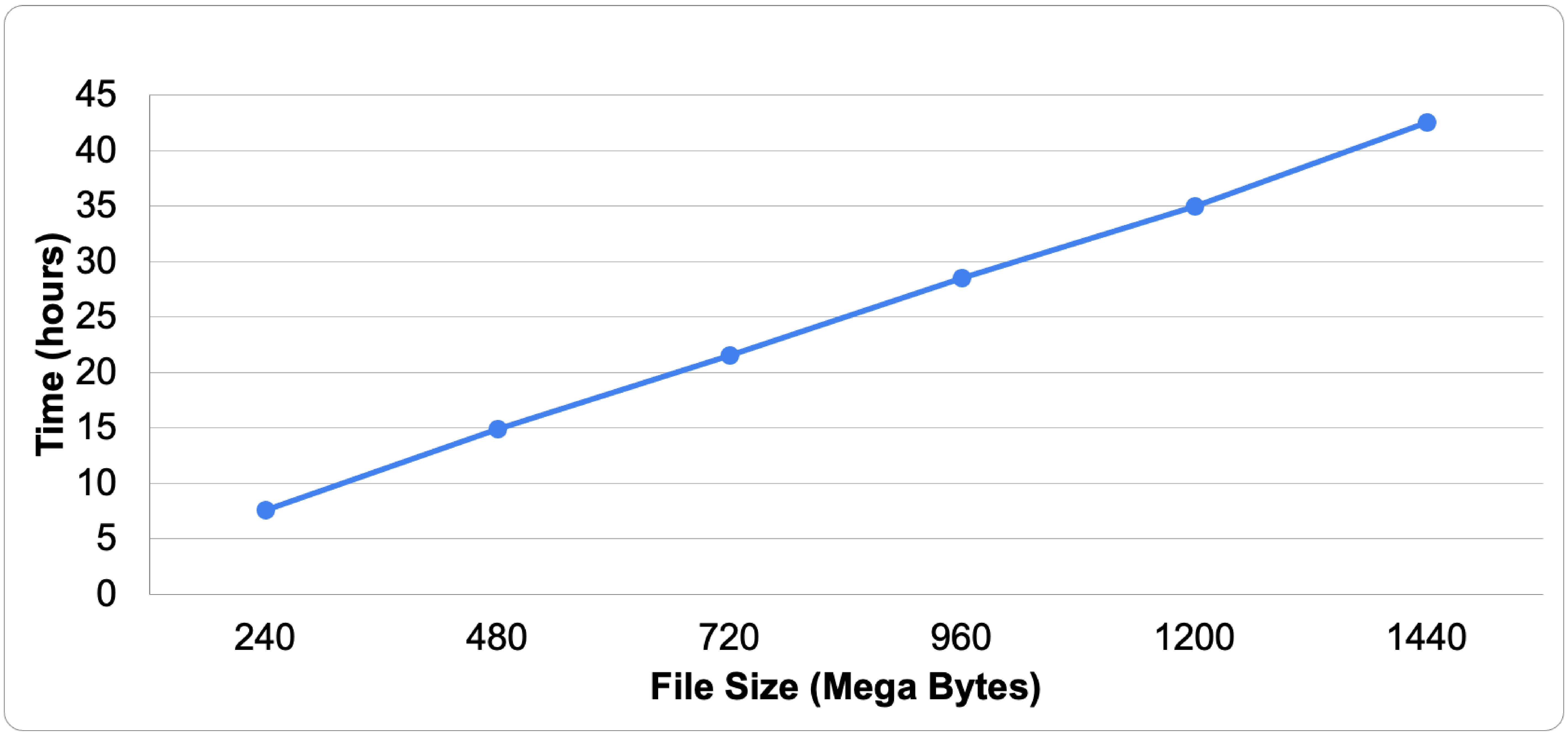


**Figure 8: Time of Input File Size vs. Hours needed to Process NBC 9mers with 180GB of RAM and 48 cores.**

**Kraken2 Varying Database**

**Micro-Recall**

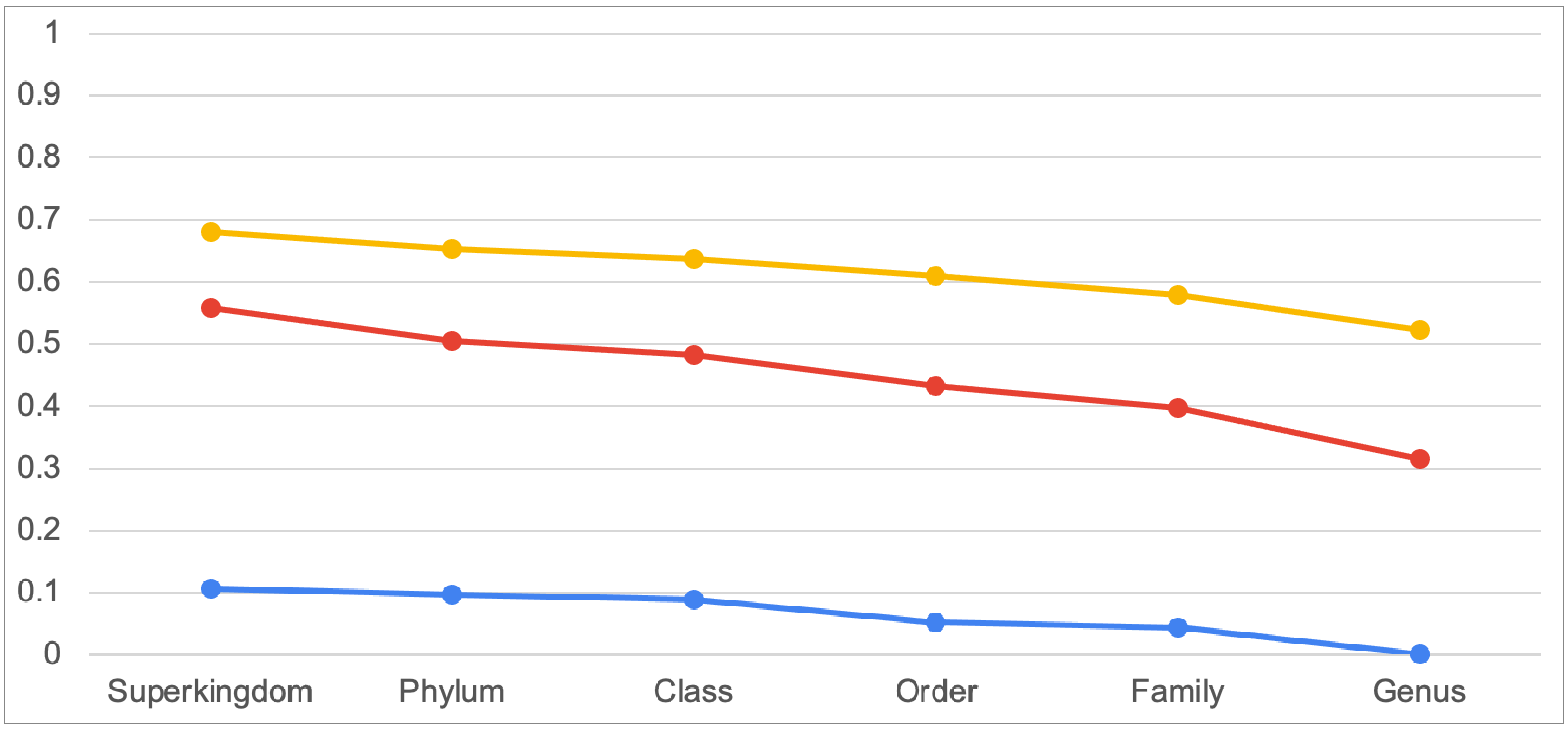


**(a)**

**Micro-Precision**

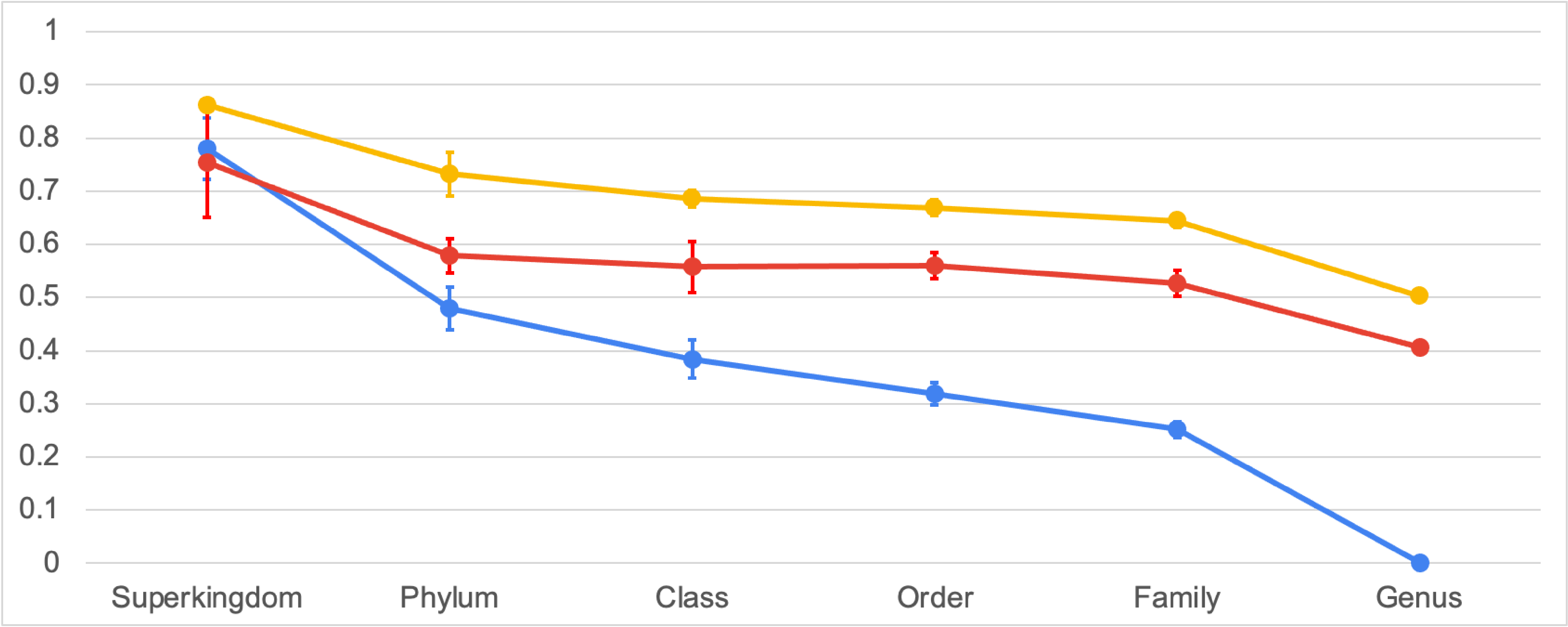


**(b)**

**Fig. 9: Kraken2's Micro-Recall/Sensitivity (a) and Micro-Precision (b) vs. Database Depth**

### Kraken2 vs. Database Depths

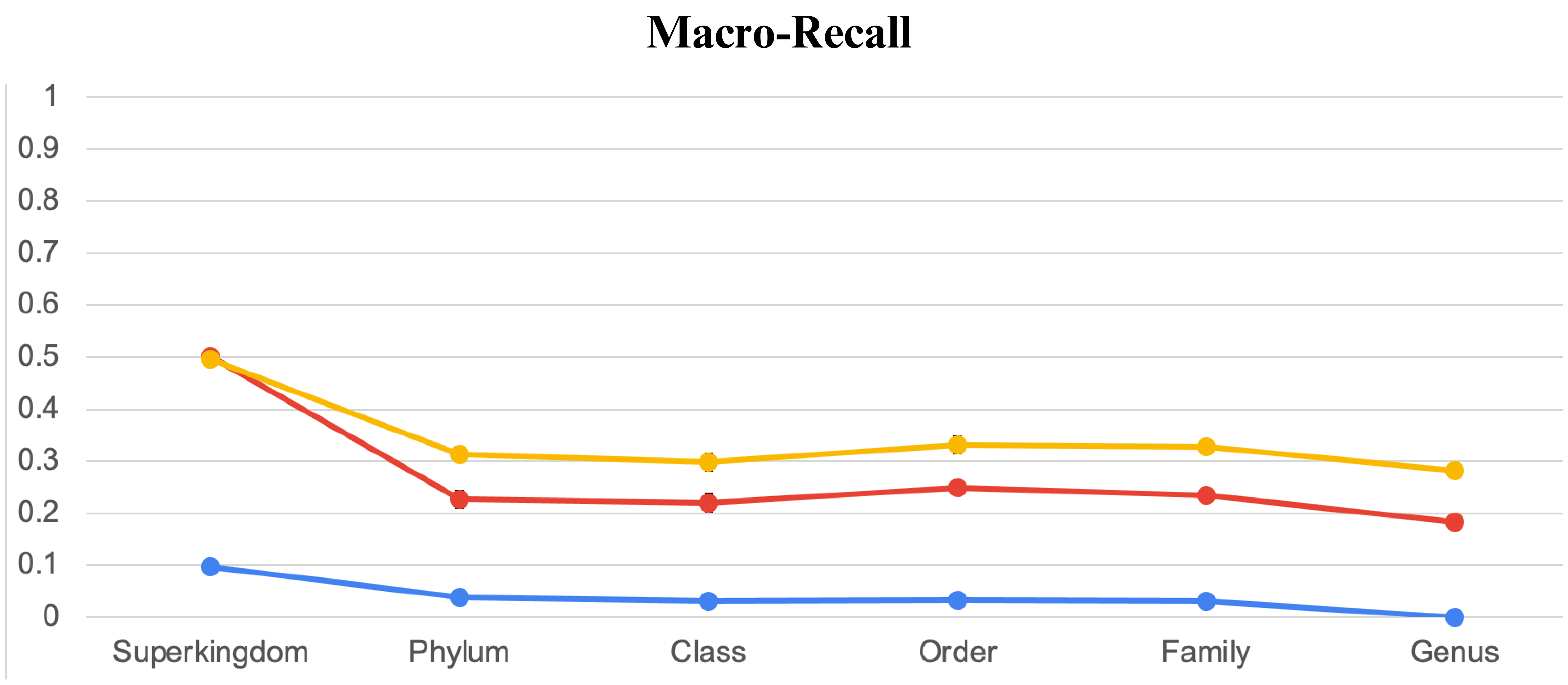


**(c)**

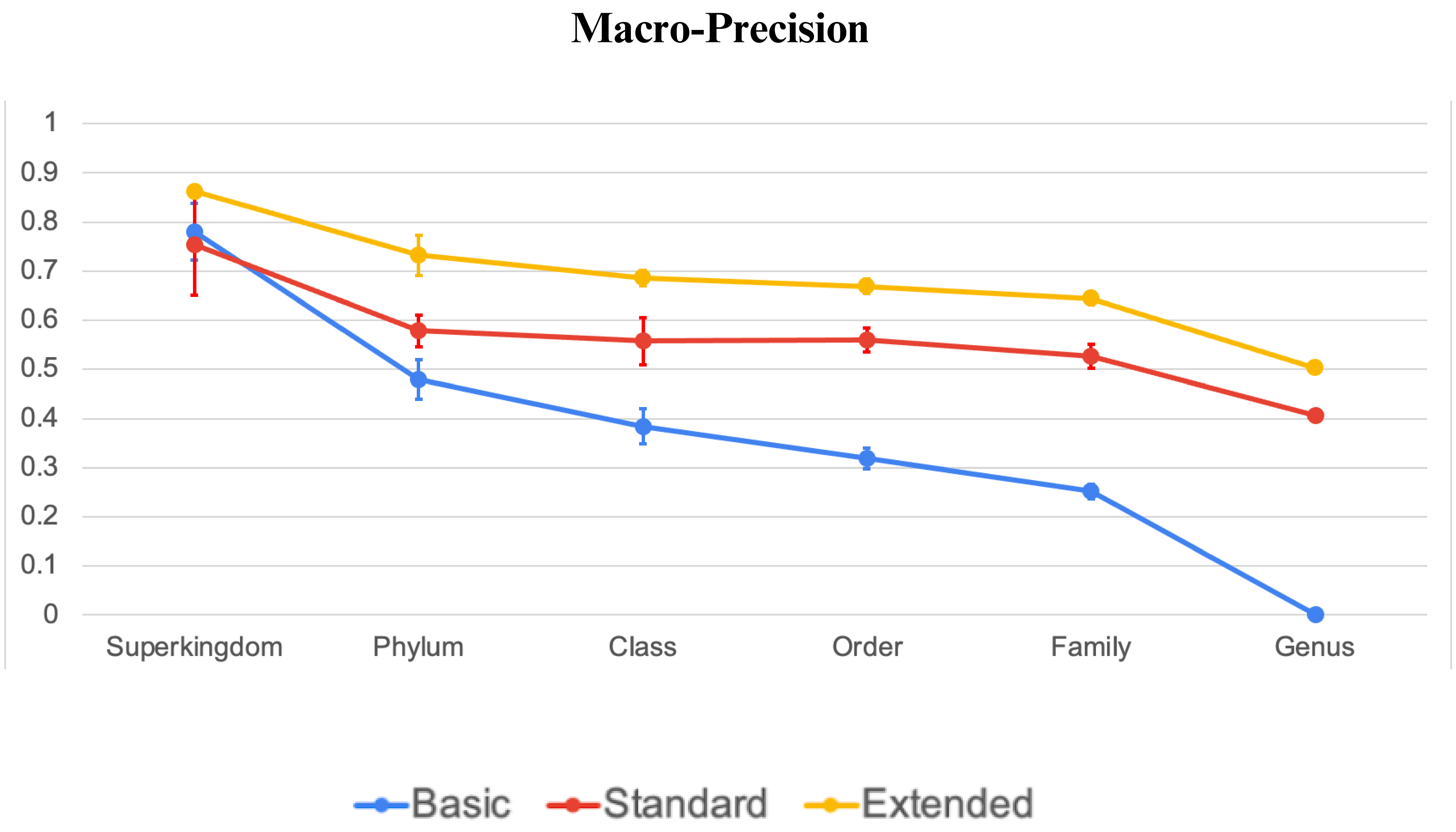


**(d)**

**Fig. 9: Kraken2's Macro-Recall/Sensitivity (c) and Macro-Precision (d) vs. Database Depths**

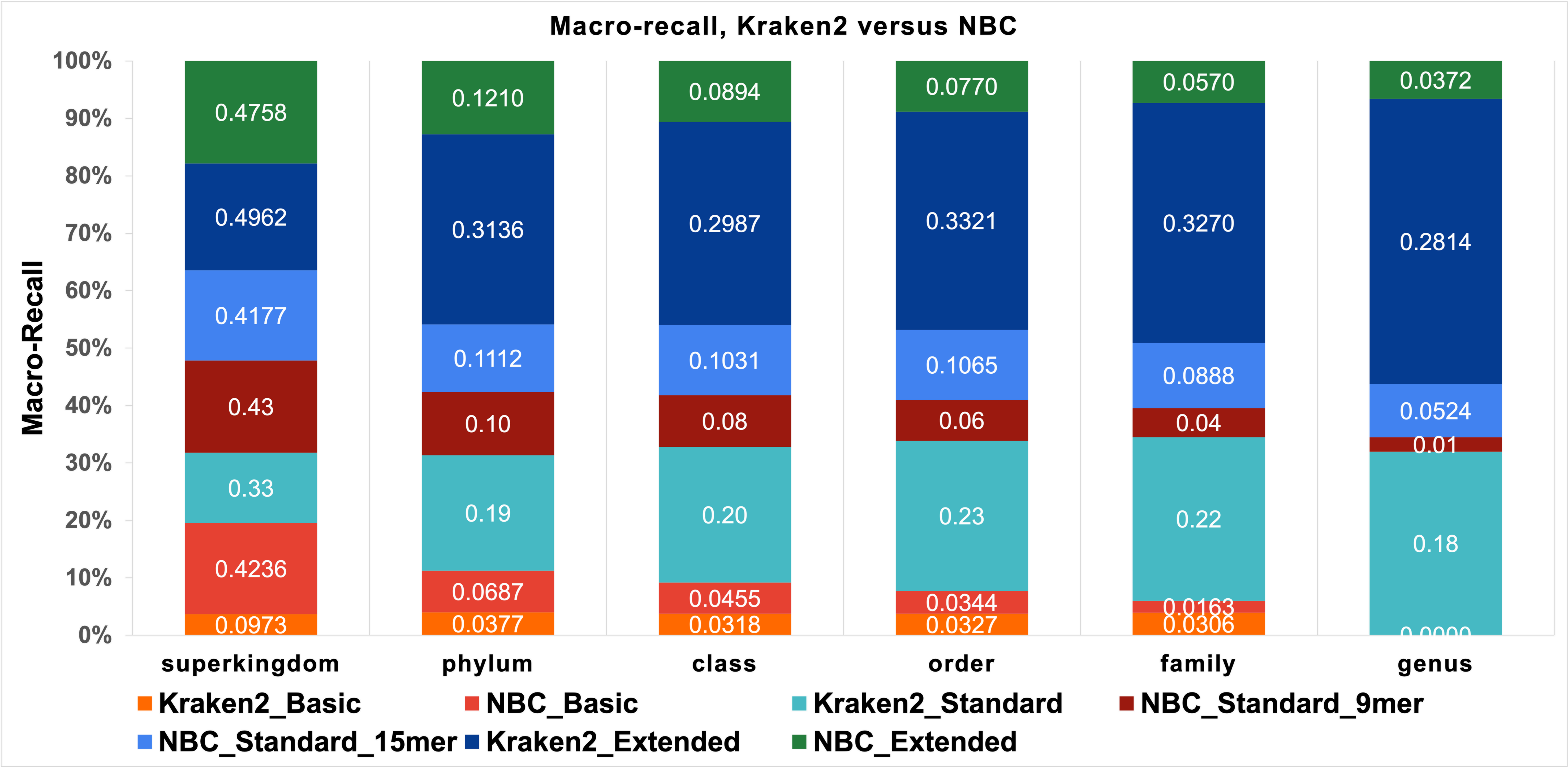


(a)

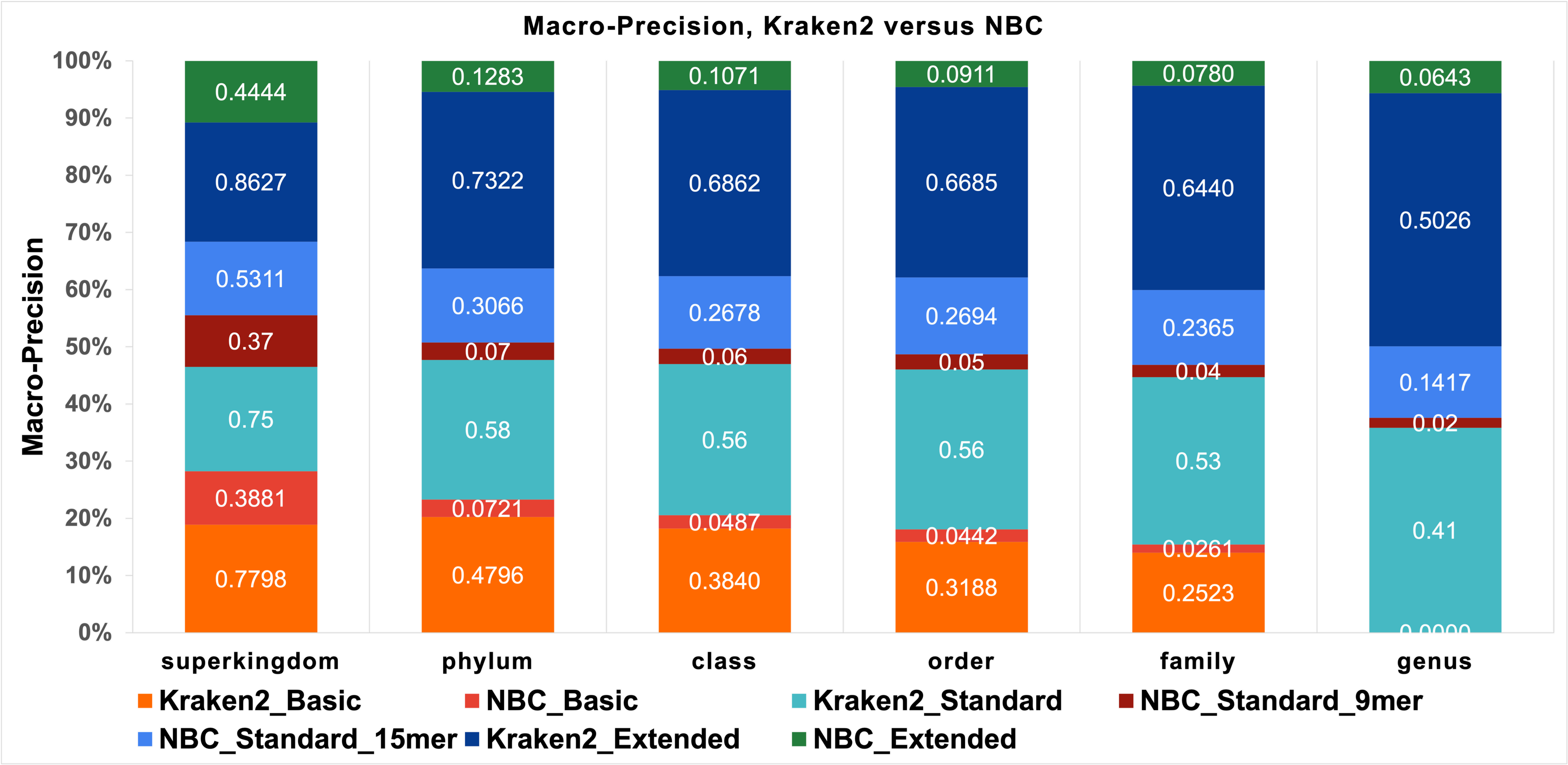


(b)

**Fig. 10: Macro-Recall/Sensitivity (a) and Macro-Precision (b) NBC vs. Kraken2 vs. Database Depth. NBC performs competitively in terms of recall for the superkingdom level, but Kraken2 calls taxa with higher precision due to the confidence threshold.**

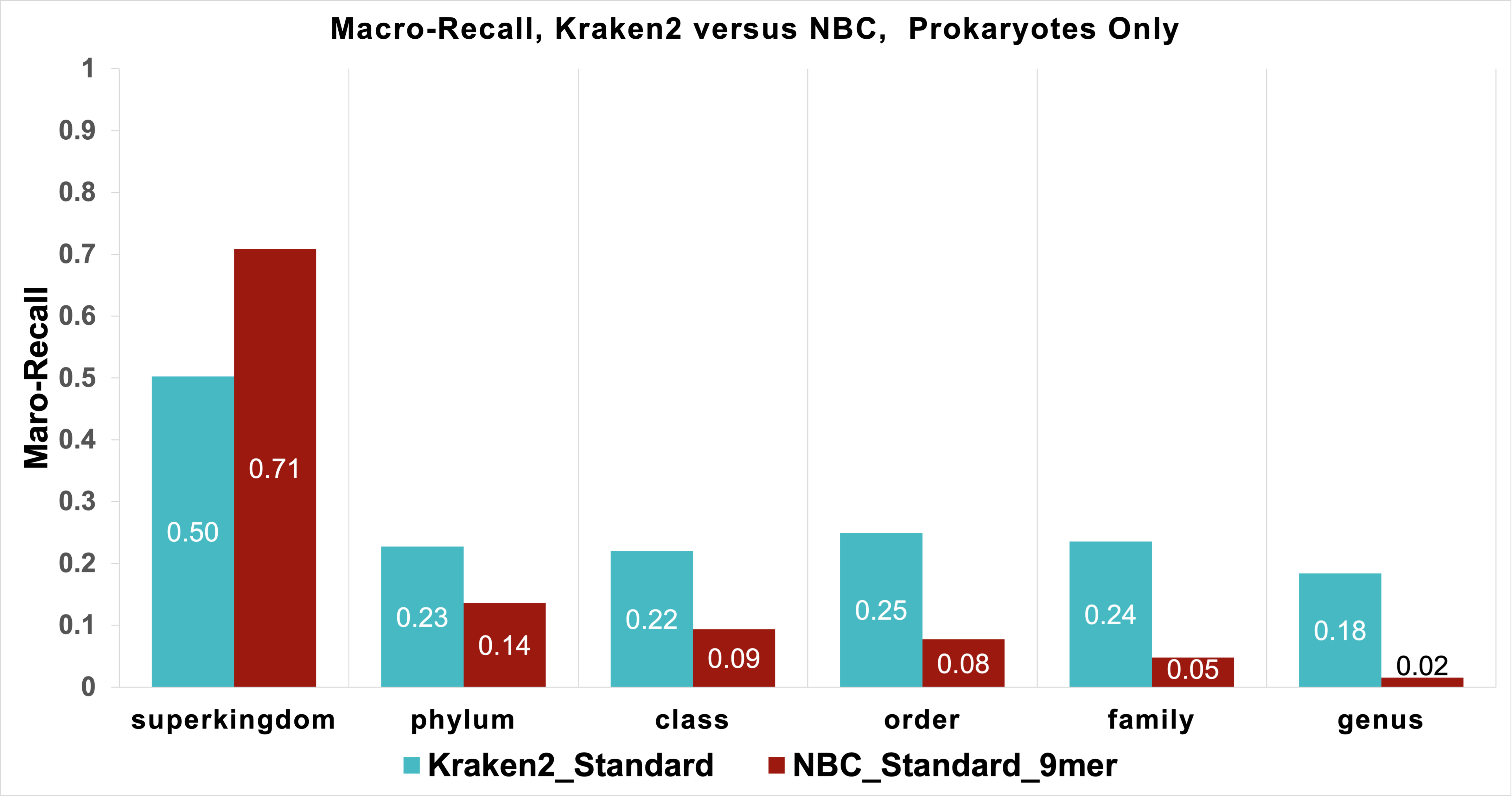


**(c)**

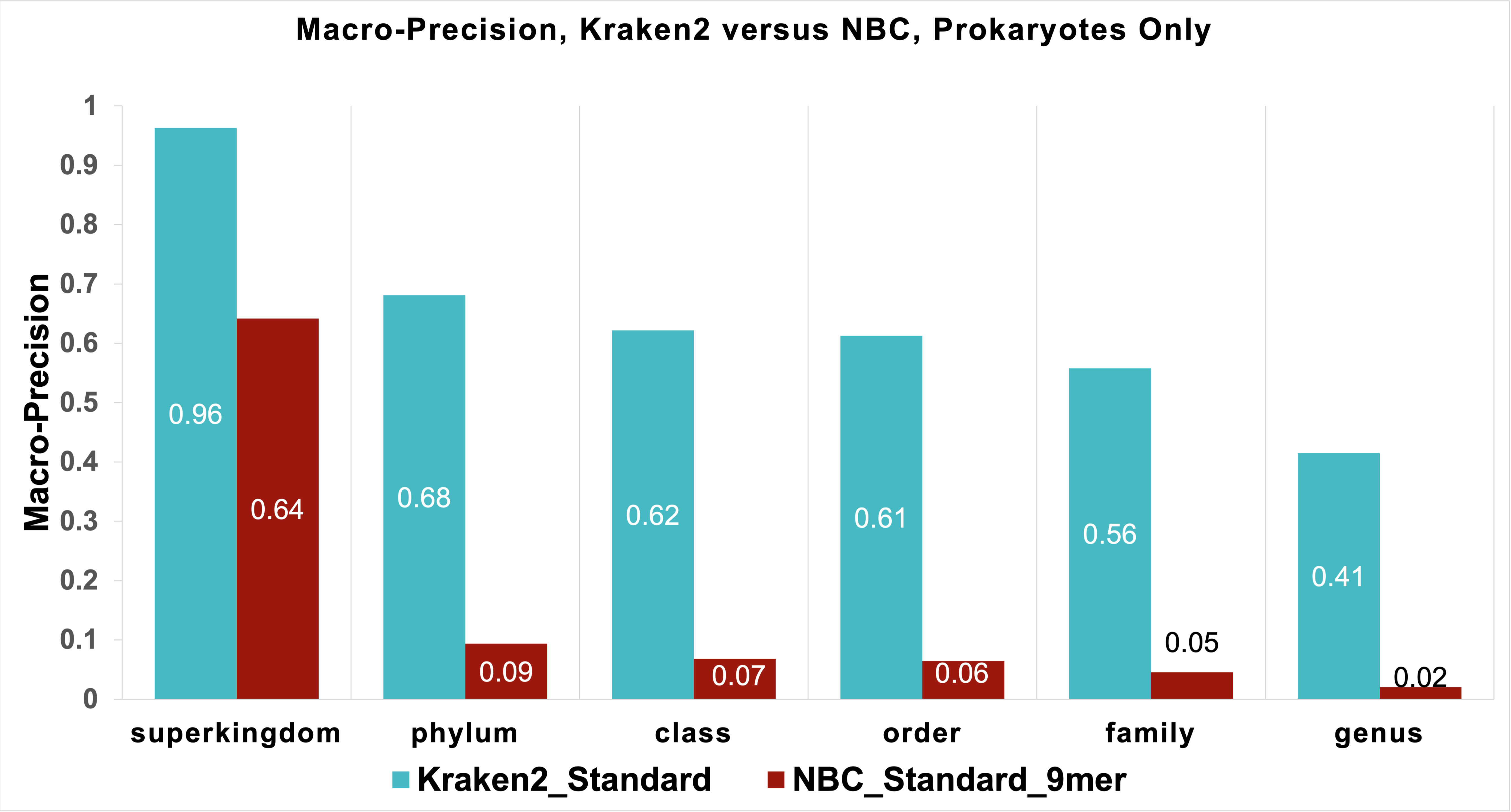


**(d)**

**Fig. 10: Macro-Recall/Sensitivity (c) and Macro-Precision (d) NBC vs. Kraken2 vs. Database Depths Prokaryotes Only**

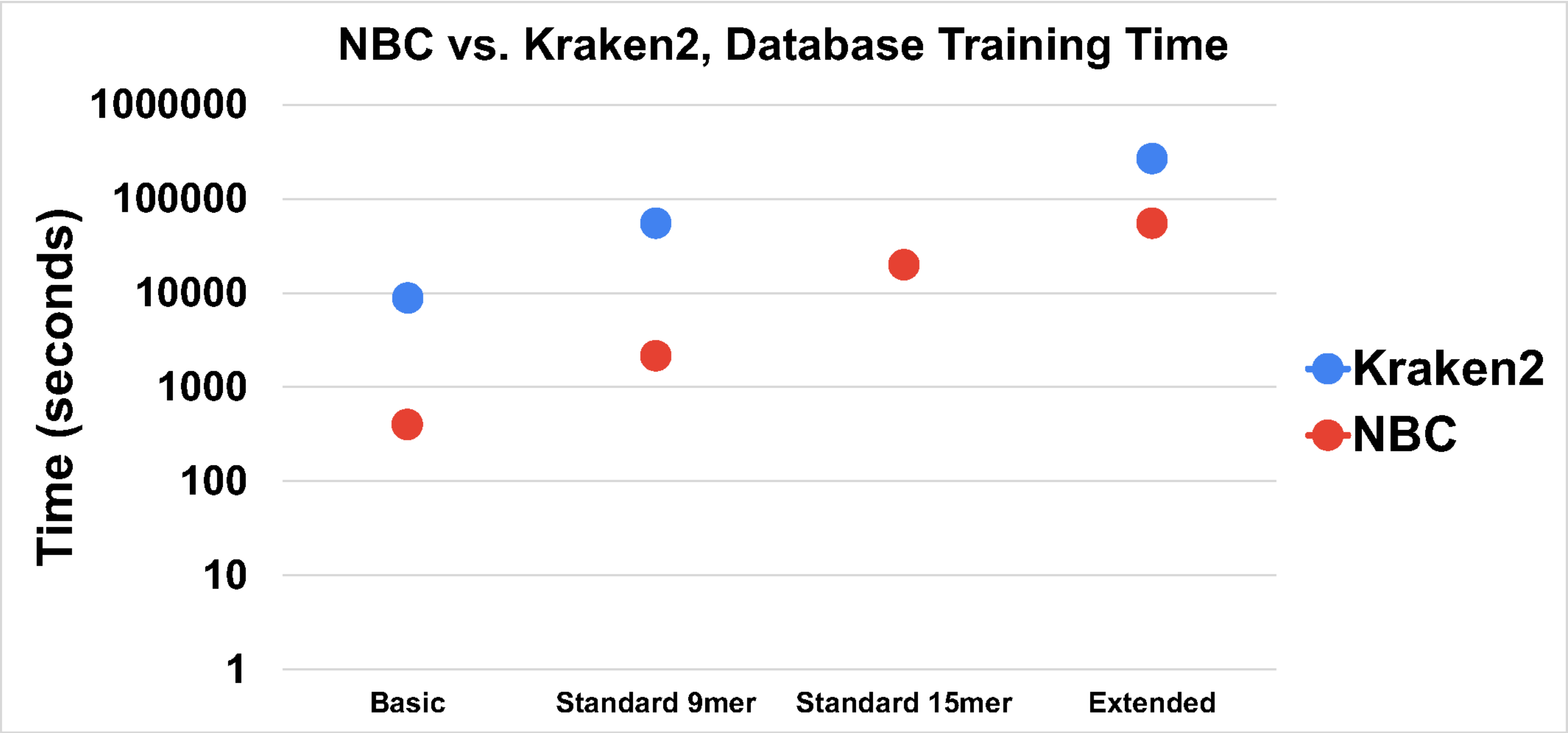


(a)

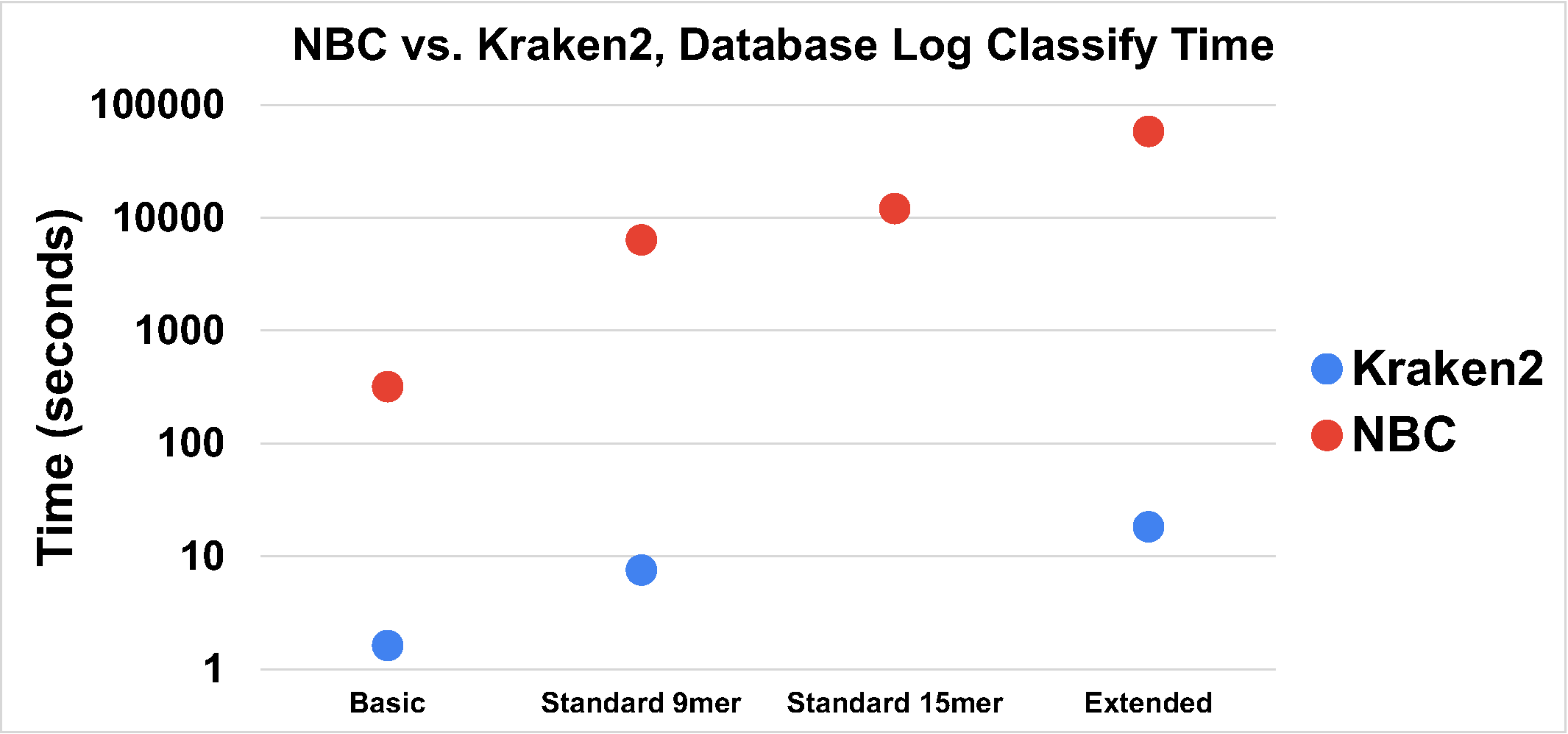


(b)

**Fig. 11 Graph of the Training (a) and Testing (b) CPU times of NBC-9mers/15mers and Kraken2 (if not stated, 9mers are used for NBC).**

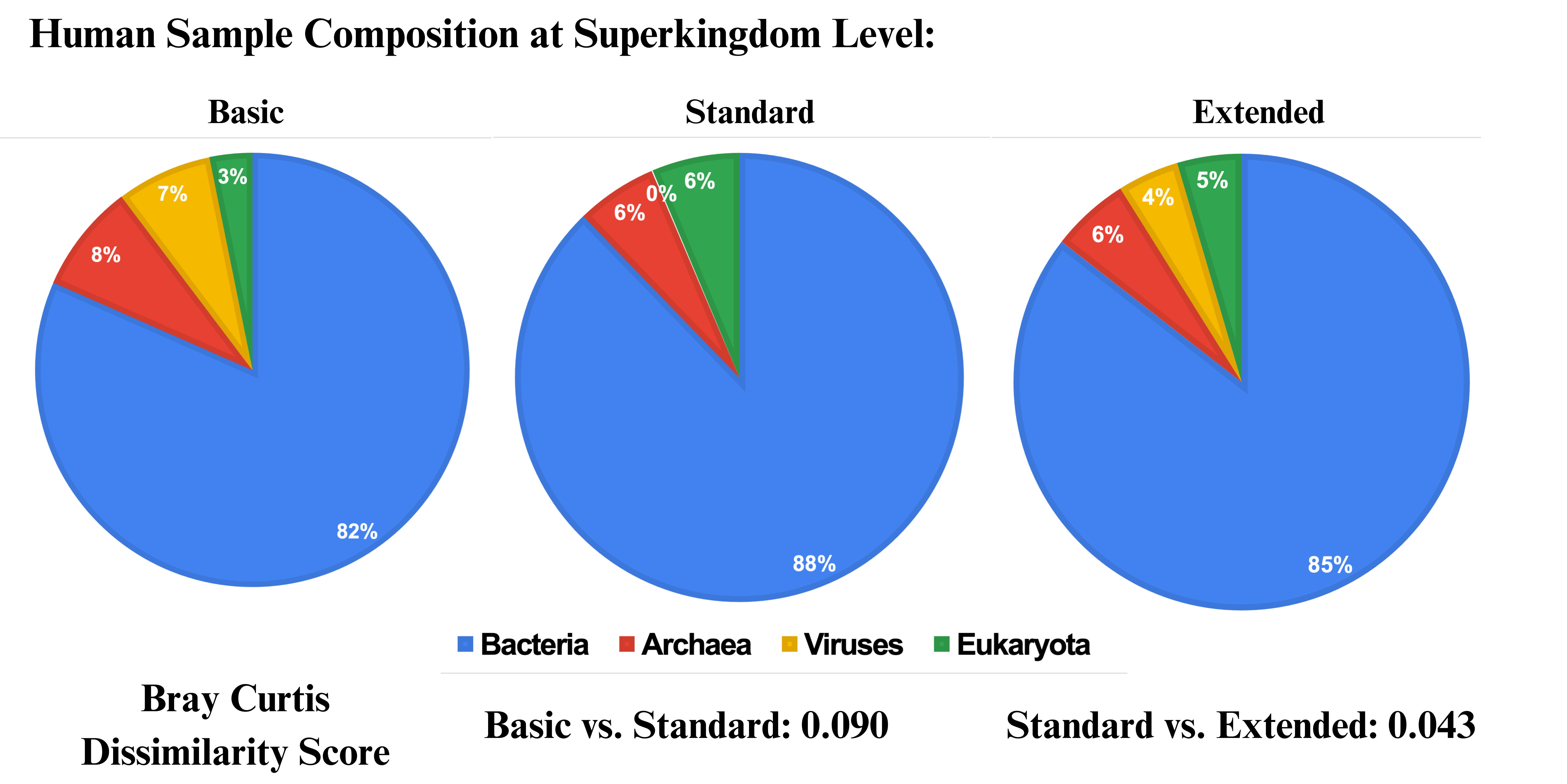


**Figure 12a: Superkingdoms Classified in the Human Gut Sample vs. Database Depth. The Standard Database has little viral representation while the Basic d**

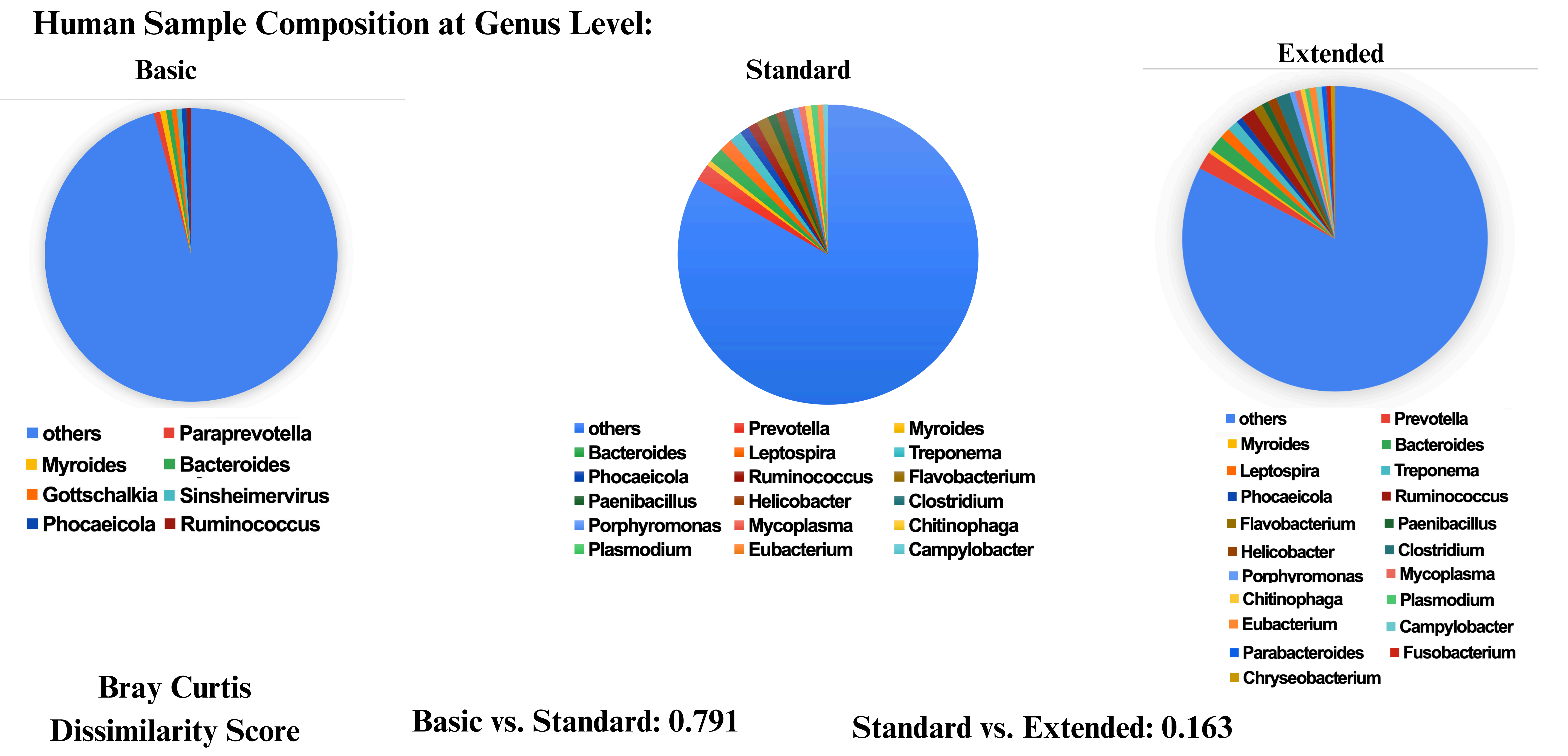


**Figure 12b: Genera Classified n the Human Gut Sample vs. Database Depth. The Basic database (which only has around one representative genome per genera is lacking major composition while the Standard and Extended are quite similar.**
